# Distinct descending motor cortex pathways and their roles in movement

**DOI:** 10.1101/229260

**Authors:** Michael N. Economo, Sarada Viswanathan, Bosiljka Tasic, Erhan Bas, Johan Winnubst, Vilas Menon, Lucas T. Graybuck, Thuc Nghi Nguyen, Lihua Wang, Charles R. Gerfen, Jayaram Chandrashekar, Hongkui Zeng, Loren L. Looger, Karel Svoboda

## Abstract

Activity in motor cortex predicts specific movements, seconds before they are initiated. This preparatory activity has been observed in L5 descending ‘pyramidal tract’ (PT) neurons. A key question is how preparatory activity can be maintained without causing movement, and how preparatory activity is eventually converted to a motor command to trigger appropriate movements. We used single cell transcriptional profiling and axonal reconstructions to identify two types of PT neuron. Both types share projections to multiple targets in the basal ganglia and brainstem. One type projects to thalamic regions that connect back to motor cortex. In a delayed-response task, these neurons produced early preparatory activity that persisted until the movement. The second type projects to motor centers in the medulla and produced late preparatory activity and motor commands. These results indicate that two motor cortex output neurons are specialized for distinct roles in motor control.

## INTRODUCTION

Motor cortex plays critical roles in planning and executing voluntary movements. Activity in motor cortex anticipates specific future movements, often seconds before movement onset (reviewed in Refs [1,2]). This dynamic neural process, referred to as preparatory activity, is thought to move the state of the motor cortex to an initial condition appropriate for eliciting rapid, accurate movements^3^. In addition, motor cortex activity is highly modulated during movement onset, consistent with commands that control the timing and direction of movements^4,5^.

Reconciling the dual roles of motor cortex requires an understanding of the cell types that make up the cortical circuit, and how these cell types integrate into the multi-regional circuits that maintain short-term memories and produce voluntary movements. Motor cortex comprises distinct cell types that differ in their location, gene expression pattern, electrophysiology, and connectivity. Intratelencephalic (IT) neurons in layers (L) 2-6 receive diverse input from other cortical areas and excite pyramidal tract (PT) neurons^6–8^. PT neurons, whose somata define neocortical L5b^9^, are of particular significance as they make the only long-range connections linking the motor cortex with premotor centers in the brainstem and spinal cord^10^. PT neurons thus coordinate cortical and subcortical brain regions to produce behavior^11,12^. Lesioning PT axons causes persistent motor deficits^12,13^. PT neurons also constitute a major component of the cortical projection to the thalamus^14–16^. Previous studies have shown that preparatory activity is not maintained by motor cortex in isolation, instead requiring reverberations in a thalamocortical loop^17^. Consistent with roles in both movement planning and initiation, PT neurons show diverse activity patterns, including preparatory activity and movement commands^18–21^. PT neurons are also structurally heterogeneous, with complex projection patterns in the midbrain and hindbrain^14^.

Here we show that PT neurons in mouse motor cortex comprise two cell types with distinct gene expression profiles and projection patterns. We refer to these cell types as PT^upper^ and PT^lower^ neurons, reflecting their distributions in different sublaminae in L5b. PT^upper^ project to the thalamus, which forms a feedback loop with motor cortex. PT^lower^ neurons project to premotor centers in the medulla. Cell type-specific extracellular recordings in the anterior lateral motor cortex (ALM) during a delayed-response task suggest that PT^upper^ neurons are involved in motor planning, whereas PT^lower^ neurons play roles in movement execution. Thus, motor cortex coordinates its two complementary roles at the level of distinct cell types.

## RESULTS

### Two types of PT neurons in Layer 5

Single-cell RNA sequencing (scRNA-Seq) was used to produce a taxonomic classification of cell types^22^ in the anterior lateral motor cortex (ALM) and in primary visual cortex (V1). A total of 21,749 scRNA-Seq transcriptomes were collected, including 9,035 from ALM. Dimensionality reduction was used to extract features from single-cell transcriptomes^23^, which in turn were the basis for clustering^22^. This procedure identified 116 transcriptomic clusters. GABAergic neurons partitioned into 49 clusters, all of which contained neurons from both ALM and V1^22^. ALM glutamatergic neurons belonged to 21 clusters (**Fig. 1**), which were distinct from the glutamatergic clusters identified in V1.

**Figure 1.**
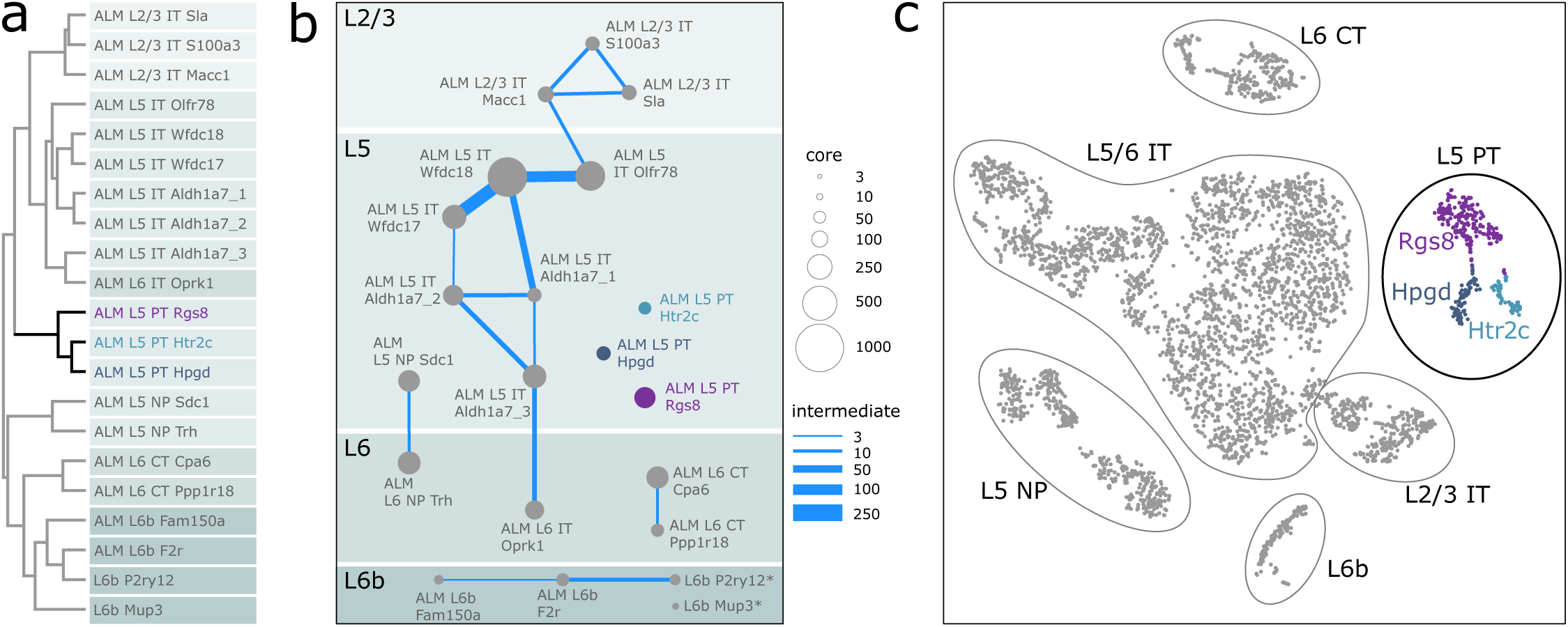
Taxonomy of motor cortex glutamatergic neurons based on single-cell RNA-seq. **a.**Hierarchical clustering of gene expression. Three gene expression clusters, identified by the genes *Rgs8*, *Htr2c,* and *Hpgd,* correspond to pyramidal tract neurons. **b.** Constellation diagram of gene expression clusters in ALM. Node diameters indicate the number of neurons belonging to each cluster (core) and edges represent cells shared by two clusters (intermediates). **c.** Two-dimensional stochastic neighbor embedding (tSNE) projection of transcriptomic data of sequenced single neurons in ALM. The cluster memberships of individual neurons are color-coded as in panels (a) and (b).

ALM and visual cortex perform different computations. Visual cortex and other sensory cortical areas process sensory information with millisecond time-scale dynamics^24,25^. ALM shows slow dynamics related to short-term memory and motor planning, in addition to fast dynamics related to the initiation of orofacial movements^18,26^. To gain an understanding of the neural circuit specializations underlying ALM function, we analyzed ALM projection neurons. Transcriptomically, ALM glutamatergic projection neurons within the same cortical layer and/or belonging to the same projection types were generally more similar to each other. Expression clusters corresponding to L2/3 intratelencephalic (IT), L5/6 IT, L5 pyramidal tract (PT), L6 corticothalamic (CT), and L6b subplate neurons exhibited a higher degree of similarity within a projection type than between types (**Fig. 1a,c**), as did L5 neurons that lack long-range projections (NP; ‘near projecting’).

PT neurons form the sole cortical projection to motor areas in the midbrain and hindbrain, and therefore likely play important roles in motor planning and execution. For the rest of this study, we focus on these neurons. ALM PT neurons, retrogradely labeled from a diverse set of PT targets^22^, mapped to three distinct transcriptomic clusters: the *Rgs8* and the closely related *Hpgd* and *Htr2c* clusters (**Fig. 1** and **EDFig. 1a**). To map the structural diversity of PT neurons, we imaged and reconstructed the brain-wide axonal projections of entire PT neurons, labeled randomly by viral injection in ALM^27^ (n = 12; **Fig. 2a,b** and **EDTable 1**; median axonal length: 121,037 μm, range: 80,873 - 188,105 μm; median branch points: 243, range: 144 - 540). Single-neuron reconstructions suggested two classes of PT neurons based on their axonal projections. Axons of one group innervated the thalamus (n=8; **Fig. 2a,b**; *yellow-green hues* and **EDFig. 2**). The other group bypassed the thalamus and branched extensively in the reticular nuclei of the medulla (n=4; **Fig. 2a,b**; *red-brown hues* and **EDFig. 2**).

**Figure 2.**
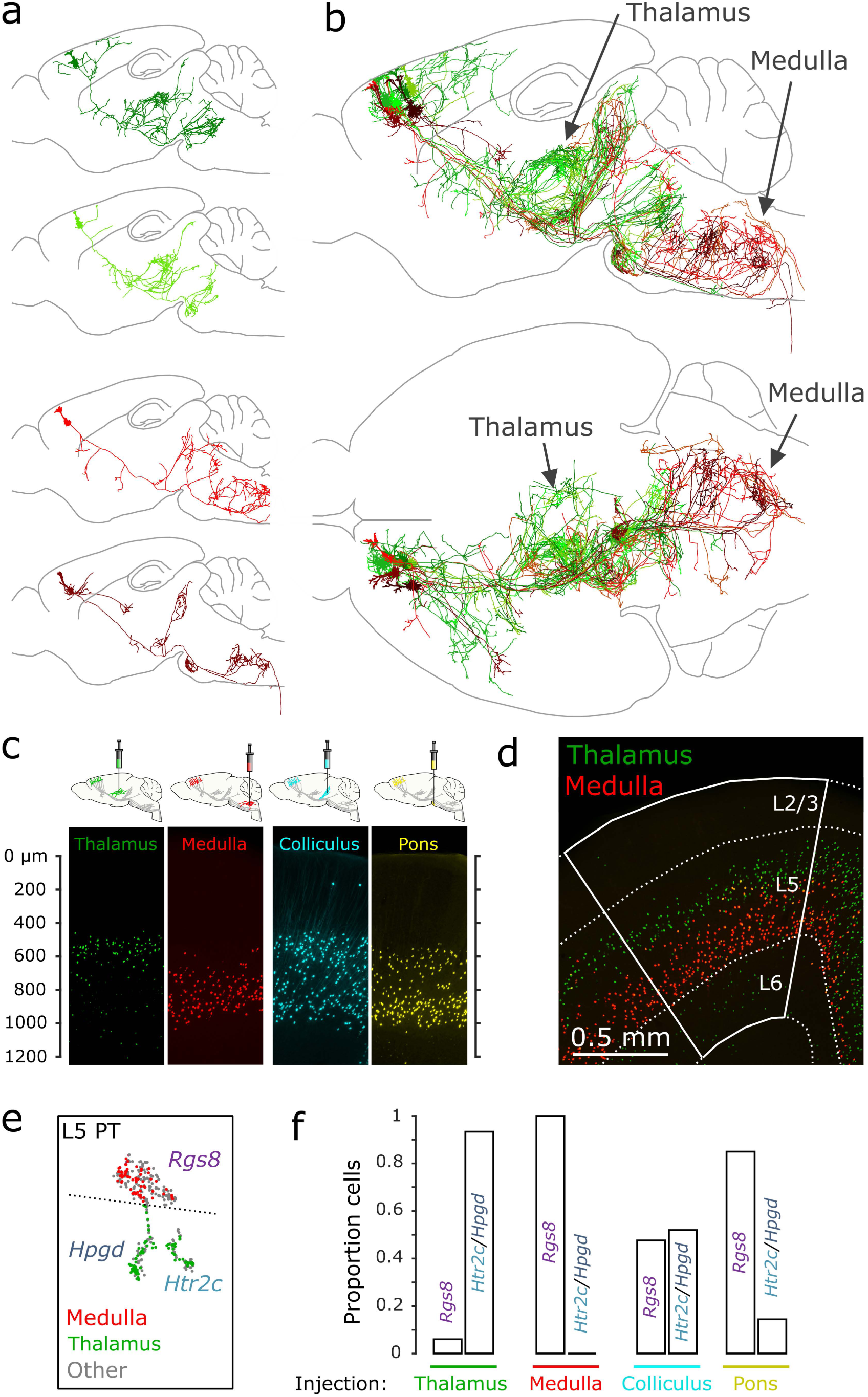
Two types of PT neuron in motor cortex. **a.** Example dendritic and axonal reconstructions of four single PT neurons. Two PT neurons project to the thalamus (*top; yellow-green hues*) and two project to the medulla (*bottom; red-brown hues*). **b.** Reconstructions of 4 thalamus-projecting PT neurons and 4 medulla-projecting PT neurons overlaid and collapsed in the sagittal (top) and horizontal (bottom) planes. Dendrites are denoted by thicker line segments. **c.** Nuclei of neurons retrogradely labeled from different PT targets are located in distinct sublaminae of L5b. **d**. Nuclei of PT neurons retrogradely labeled from the thalamus (*green*) or the medulla (*red*). **e.** Gene expression of PT neurons retrogradely labeled from the thalamus (*green*) and medulla (*red*) in tSNE space (as in Fig. 1c). PT neurons projecting to the medulla belong to the *Rgs8* cluster, whereas thalamus-projecting PT neurons were part of the *Htr2c* and *Hpgd* clusters. **f.** Proportion of neurons retrogradely labeled from each PT target that were clustered into the *Rgs8* and *Htr2c/Hpgd* expression clusters.

To determine the spatial distribution of these projection types in motor cortex, neurons were retrogradely labeled from the thalamus and medulla using AAVretro^28^. Thalamus-projecting PT neurons were located in upper L5b (**Fig. 2c,d** and **EDFig. 4**; *green cells*) and medulla-projecting PT neurons in lower L5b (*red cells*). This pattern was similar across all of motor cortex, including both primary and secondary motor areas. Retrograde labeling from the superior colliculus (SC) and pons labeled PT neurons across both L5b sublaminae, with pons-projecting neurons concentrated somewhat in the deeper sublayer (**Fig. 2c**). Consistent with their laminar distributions, few neurons (2.2%; 22/984) were co-labeled by injections into the thalamus and medulla. This lack of overlap did not result from inefficient retrograde labeling; in experiments in which neurons were retrogradely labeled from both the thalamus and SC, most PT neurons (77.1%; 687/890) projecting to the thalamus were co-labeled.

To link projection classes and transcriptomic clusters, we examined gene expression in medulla-projecting and thalamus-projecting PT neurons, isolated from AAVretro-labeled brains (Methods). All (63/63) medulla-projecting PT neurons mapped to the *Rgs8* taxonomic cluster. Similarly, thalamus-projecting PT neurons mapped to the *Htr2c* and *Hpgd* clusters (88/94; *Htr2c*, 33/94; *Hpgd*, 55/94) (**Fig.1c, 2e,f**). Furthermore, all thalamus-projecting and medulla-projecting PT neurons (157/157) could be separated using a single linear boundary in t-SNE space (**Fig. 2e**; *dotted line*). PT neurons retrogradely labeled from the pons and SC mapped to the same PT clusters (SC: *n*=92 total; 44 *Rgs8*, 20 *Htr2c*, 28 *Hpgd*; pons: *n*=100 total; 88 *Rgs8*, 9 *Htr2c*, 3 *Hpgd*; **Fig. 2f**).

Axonal reconstructions and transcriptomic data suggest that PT neurons can be divided into two distinct cell types in motor cortex. To determine if thalamus-projecting and medulla-projecting PT neurons account for the majority of PT neurons in motor cortex, we examined additional PT neurons that were reconstructed partially (thalamus-projecting: n=3; medulla-projecting: n=3). All (18/18) partially and fully reconstructed PT neurons projected to the SC and no (0/18) PT neurons lacked projections to both the thalamus and medulla (**EDTable 1**). These results suggest that most – if not all – PT neurons are accounted for by the medulla-projecting and thalamus-projecting types. We refer to the superficial, thalamus-projecting *Htr2c/Hpgd* cell type as PT^upper^ neurons and the deep, medulla-projecting *Rgs8* cell type as PT^lower^ neurons, reflecting their laminar distributions.

### Cell type-specific markers

We combined scRNA-Seq with bulk RNA-Seq to identify marker genes for PT^upper^ and PT^lower^ cells. Bulk RNA-Seq data was collected for PT^upper^ and PT^lower^ neurons from AAVretro-labeled brains (50-70 cells/sample; 6 replicates each; **Methods**). Expression levels in scRNA-Seq and bulk RNA-Seq were highly correlated (Pearson’s R = 0.86-0.88; **EDFig. 5**). Approximately 11,000 genes were detected per cell type in bulk RNA-Seq, compared to approximately 10,000 in the scRNA-Seq data (median of detected genes for single cells: PT^upper^ 9936 genes; PT^lower^ 9865 genes). Differentially expressed (DE) genes identified from scRNA-Seq were also differentially expressed in bulk RNA-Seq (**EDFig. 6a**). Conversely, DE genes from bulk RNA-Seq (**EDFig. 6b**) displayed consistent relative expression levels in scRNA-Seq (**EDFig. 1b**). Differentially expressed transcripts (**Fig. 3a**) were examined in the Allen Brain Atlas (http://mouse.brain-map.org) for enrichment in L5. Two transcripts, *Npnt* and *Slco2a1,* were confirmed as cell-type specific markers with single-molecule RNA fluorescence *in situ* hybridization (smFISH; **Fig. 3b,c**). PT^lower^ neurons expressed higher levels of *Slco2a1* mRNA (inter-quartile range, IQR = 9-28 puncta) than PT^upper^ neurons (IQR = 0-3 puncta). In contrast, PT^upper^ neurons expressed higher levels of *Npnt* (IQR = 15-30 puncta) than PT^lower^ neurons (0-4 puncta).

**Figure 3.**
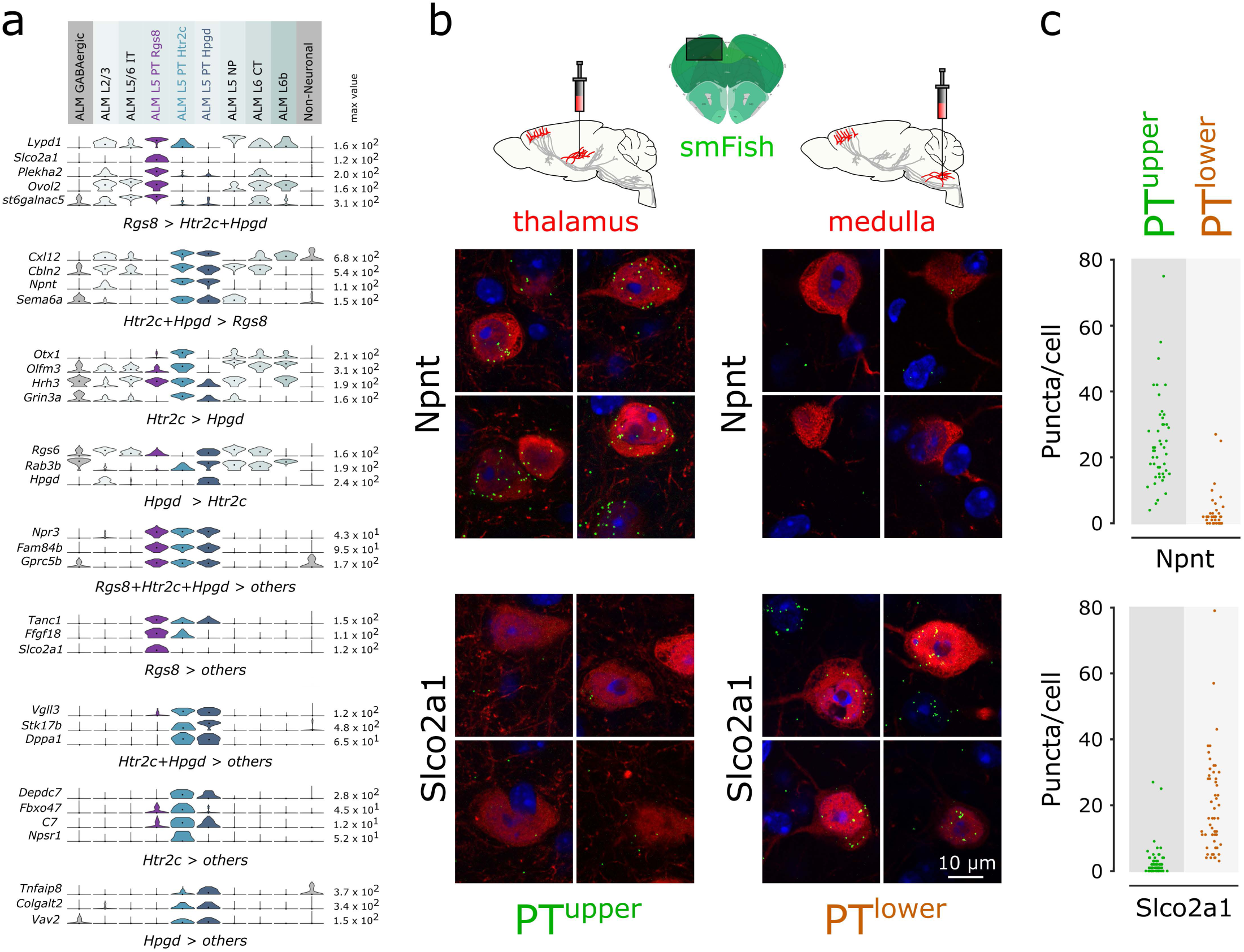
Cell type markers. **a.** Differentially expressed genes from scRNA-seq, represented by violin plots. Each row represents a single gene, and values within rows are normalized between 0 and the maximum expression value for each gene (right edge of each row; FPKM) and displayed on a log_10_ scale. Median values are shown as black dots within each violin. Differentially expressed genes are grouped by the transcriptomic clusters that they differentiate. ‘Others’ refers to ALM non-PT expression clusters. **b.** Single-molecule fluorescence *in situ* hybridization validating cell type-specific marker expression. Four example fields of view are shown for each gene and projection class. Neurons labeled from the thalamus or medulla are shown in red and RNA puncta are in green. **c.** Number of RNA puncta counted in cells of each type. Number of cells: PT^upper^: 50 *Npnt*, 66 *Slco2a1;* PT^lower^: 39 *Npnt*, 56 *Slco2a1*.

### Cell type-specific recordings

The projection patterns of the PT cell types suggest distinct roles in motor control. The cortico-thalamocortical loop is necessary for maintaining persistent preparatory activity related to motor planning^17^. PT^upper^ cells project to the thalamus and lack projections to premotor nuclei in the medulla. These characteristics suggest a role for PT^upper^ cells in generating and/or maintaining preparatory activity. In contrast, PT^lower^ cells project to premotor centers in the medulla and the spinal cord (**Fig. 2a,b** and **EDFigs. 2,3**), with few collaterals in the basal ganglia and thalamus, suggesting a role in movement execution.

We performed projection-specific recordings in ALM during a delayed-response task^20,26^ (**Fig. 4a,b**). Mice were trained to discriminate object locations with their whiskers^26^ and signal their decision about object location with skilled, directional licking (‘left / right’), but only after a delay epoch lasting 1.3 seconds. The delay epoch was terminated by an auditory ‘go’ cue instructing animals to respond. ALM is a hub for planning and executing movements in this task^18,26,29,30^.

**Figure 4.**
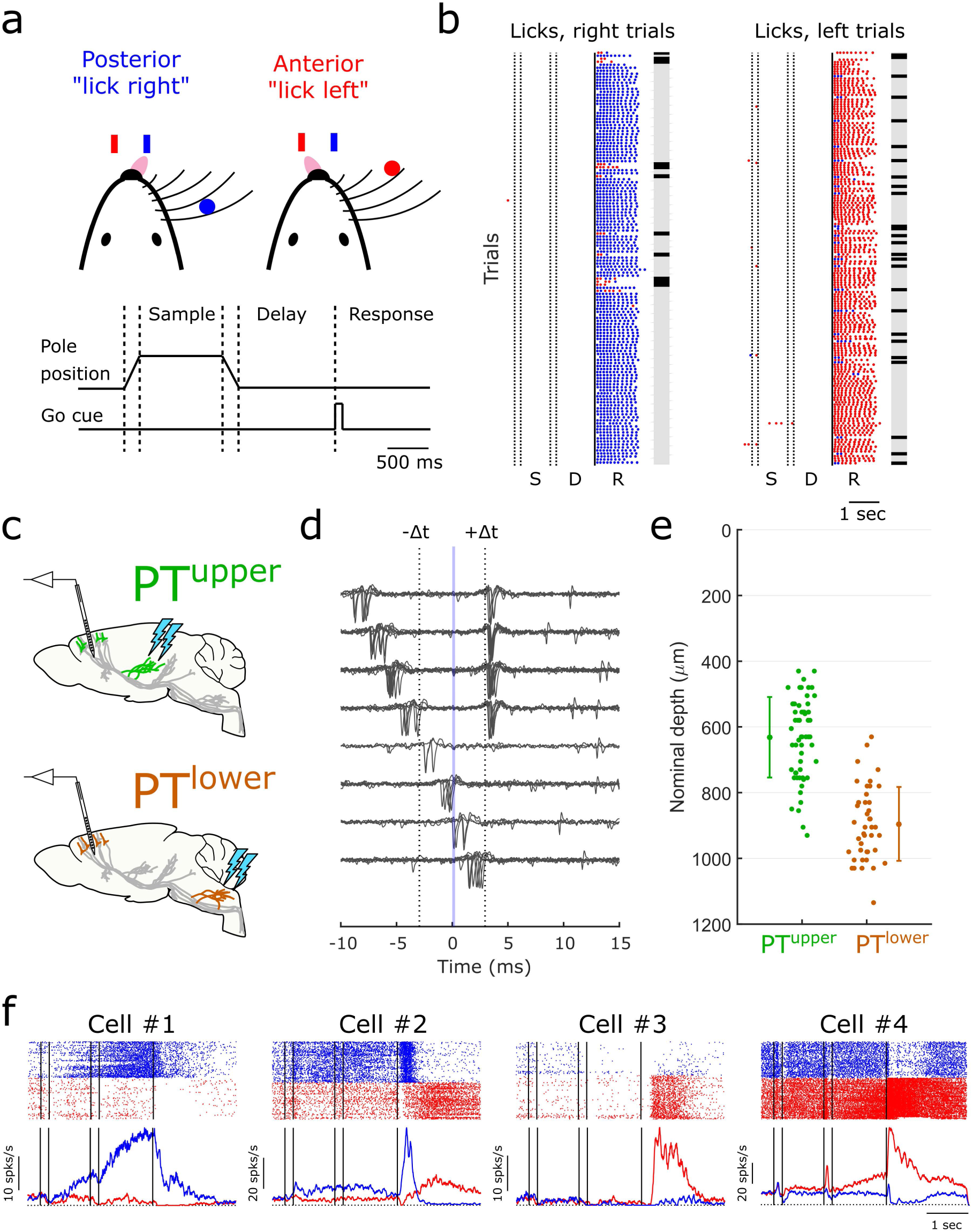
Cell type-specific extracellular neurophysiology. **a.** Mice were trained on a delayed-response task. On each trial, an object appeared within reach of the whiskers in one of two rostro-caudal positions during the sample epoch (1.0 s). The pole was removed and after a brief delay epoch (1.3 s), mice reported the pole position by licking a reward port on the right (caudal pole position) or the left (rostral pole position). **b.** Performance during an example session. Dots represent licks to the right (*blue*) or the left (*red*). Gray and black marks indicate correct and incorrect trials, respectively. **c.** Schematic for stimulation and recording configuration for each cell type. **d.** Collision test for an example neuron. Trials with spontaneous spikes preceding the light-evoked spike are shown binned by the latency of spikes preceding the stimulus from top to bottom. Putative photostimulation-evoked spikes are at +Δt. When a spike occurs in the interval [−Δt +Δt], a collision occurs with the photostimulation-evoked spike in the axon and the spike at +Δt is absent. **e.** Depth distribution of PT^upper^ and PT^lower^ neurons based on micromanipulator readings. Depths are measured from the dorsal surface and are uncorrected for the curvature of cortical layers. Error bars represent mean ± s.d. **f.** Example identified neurons. *Top:* spike rastergrams for correct lick right trials (blue) and lick left trials (red). *Bottom:* trial-averaged spike rates. S, sample; D, delay; R, response.

PT^upper^ or PT^lower^ cells were infected with AAVretro expressing channelrhodopsin-2 (ChR2). Fiber optic cannulae were implanted into the thalamus (to activate PT^upper^ cells; **Fig. 4c**, *top*) or medulla (to activate PT^lower^ cells; **Fig. 4c**, *bottom*). ChR2-expressing cells in ALM were identified with axonal photostimulation and extracellular recordings in ALM using a collision test (61 PT^upper^ cells, 8 mice; 69 PT^lower^ neurons, 4 mice; **Fig. 4d** and **EDFig. 7**)^18^. Identified PT^upper^ cells were at more superficial recording depths than PT^lower^ neurons, consistent with the retrograde labeling experiments (**Fig. 4e** and **EDFig. 4**). Layer 6 corticothalamic cells, which also innervate the thalamus, were inefficiently labeled by AAVretro (**Fig. 2c,d** and **EDFig. 4**)^28^ and excluded based on recording depth, several hundred micrometers deeper than the PT^upper^ cells. Baseline and trial-averaged peak spike rates were not significantly different (p > 0.1, two-sample t-test) in PT^upper^ cells (baseline: median=4.3 ± 3.5 Hz; peak: median=17.7 ± 13.8 Hz) and PT^lower^ cells (baseline: median=5.2 ± 4.8 Hz; peak: median=17.2 ± 22 Hz). Spike rates in PT^lower^ cells were more heterogeneous across the population (baseline: p = 0.02; peak: p = 3x10^-4^; χ^2^ test). A substantial proportion of PT^lower^ neurons displayed spike bursts (18.8%), which were rare among PT^upper^ cells (3.3%; p = 0.006, Fisher’s exact test; **EDFigs. 7,8**).

### Preparatory activity

Individual neurons exhibited diverse patterns of activity and selectivity, defined as the difference in spike rate between trial types (“lick left” vs “lick right”) (**Fig. 4f**). Most recorded PT neurons had significant selectivity (p<0.01, two-sided Mann-Whitney U-test) during at least one task epoch (122/130; 94%). A subset displayed selectivity that emerged at the start of the sample epoch and persisted through the delay epoch until the response epoch (**Figs. 4f**; *left cells*), suggesting that a subset of PT neurons stably encode upcoming movement direction. This preparatory activity is a form of short-term memory that links past events and future movements.

We investigated the emergence and maintenance of preparatory activity in populations of PT^upper^ and PT^lower^ cells. We analyzed population dynamics in activity space, where each dimension corresponds to the activity of one neuron. Preparatory activity for different movement directions corresponded to distinct trajectories in the high-dimensional activity space. For each population, we computed the linear combination of cells that best discriminated trial type during the first 400 ms of the sample epoch (CD_early_: ‘early coding dimension’; **Fig. 5a,b**). We then projected the trial-averaged activity of all cells in the population along CD_early_ (**Fig. 5a**) to produce the one-dimensional projection of each activity space trajectory with maximal selectivity during this time period. In this projection, selectivity was larger and more consistent across trials in the PT^upper^ population compared to the PT^lower^ population (**Fig. 5b**). Furthermore, selectivity in the PT^upper^ population remained constant throughout the sample and delay epochs and until the go cue. We conclude that the PT^upper^ population retained decision-related information for the duration of the behavioral trial. In contrast, selectivity in the PT^lower^ population decayed rapidly along CD_early_ and was lost at the time of the go cue (no significant selectivity; p = 0.15, bootstrap). This decay did not reflect a lack of any selectivity in the PT^lower^ population; along a different dimension in activity space that maximized selectivity at the end of the delay epoch, CDlate, selectivity was substantial in both cell types (**EDFig. 9**; PT^upper^: 40/61; PT^lower^: 44/69)^18^. We computed the stability of the coding dimension across trial time. For the PT^upper^ population, the CD remained similar across the sample and delay epochs, whereas for the PT^lower^ population, the CD was uncorrelated across epochs (**EDFig. 10**). As suggested by population analyses, individual PT^upper^ neurons displayed persistent selectivity throughout the sample and delay epochs (**Fig. 5c**) and decoded trial type significantly better than PT^lower^ neurons during the sample epoch (**EDFig. 11**).

**Figure 5.**
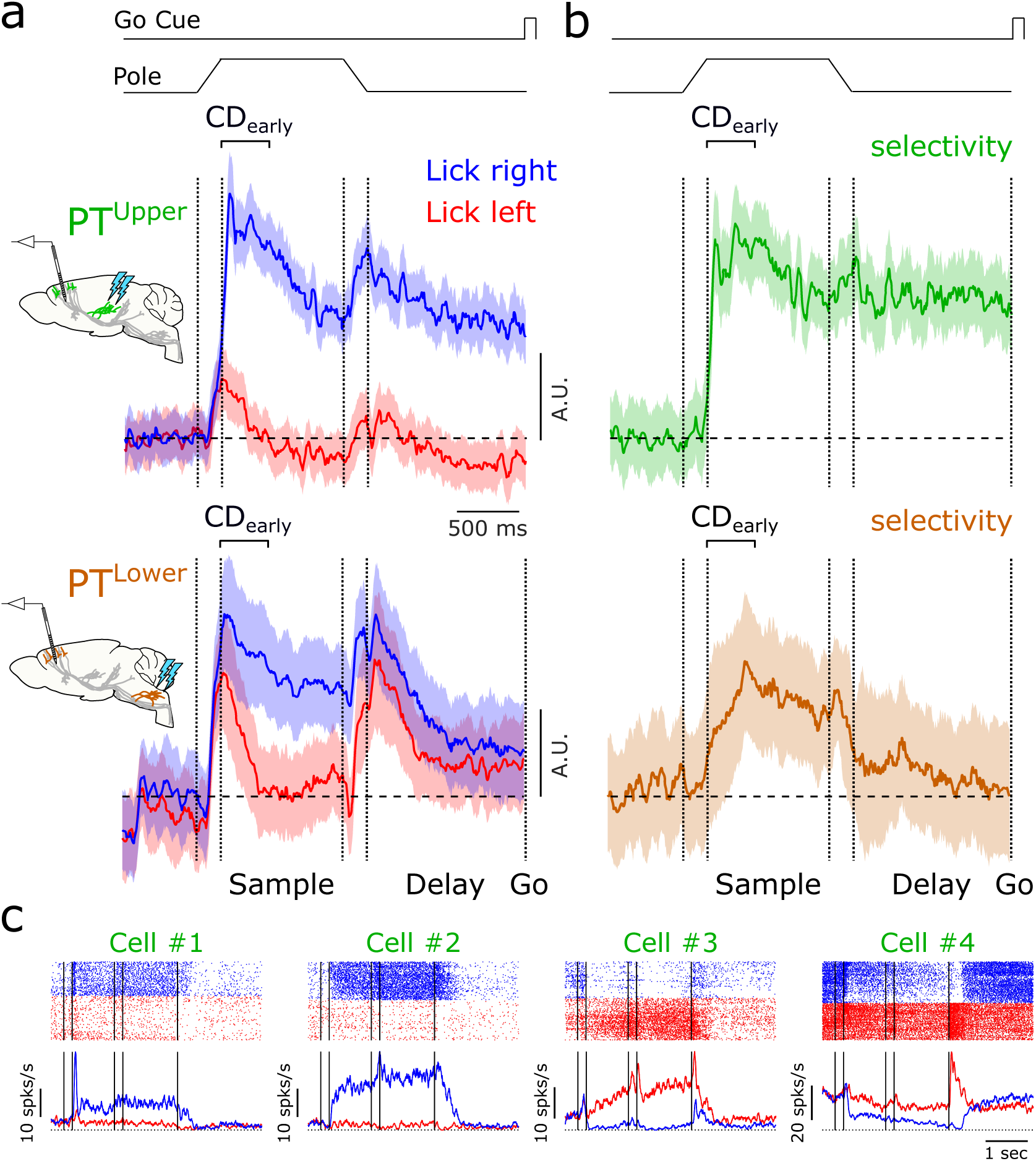
Persistent preparatory activity in PT^upper^ neurons. **a.** Time-course of the linear combination of neuronal activity that best differentiates trial type after stimulus onset (CD_early_) on lick right (*blue*) and lick left (*red*) trials for PT^upper^ (*top*) and PT^lower^ (*bottom*) neurons. **b.** Difference in CD_early_ projections on lick right and lick left trials (selectivity) in PT^upper^ (*top; green*) and PT^lower^ (*bottom; orange*) neurons. **c.** Example identified PT^upper^ neurons. *Top:* raster plots for correct lick right trials (blue) and lick left trials (red). *Bottom:* trial-averaged spike rates. Shaded regions in (a) and (b) represent 95% confidence intervals around the mean using hierarchical bootstrapping.

### Movement commands

The rhythmic movements involved in licking and swallowing are coordinated by circuits in the reticular nuclei of the medulla^31,32^. Microstimulation of ALM is sufficient to initiate directional licking^18,33^. PT^lower^ neurons provide a direct path from ALM to the premotor circuits in the medulla (**Fig. 2a,b** and **EDFigs. 2,3**). We reasoned that putative ALM signals driving movement should have selectivity for movement and emerge before movement onset. In addition, movement commands should lie along a dimension in activity space that is orthogonal to the dimension that predicts movement in the delay epoch; otherwise movement would be triggered before the go cue^34,35^.

For each population, we determined CD_go_ as the dimension that maximizes selectivity immediately after the go cue (400 ms), orthogonal to CD_late_ (**Fig. 6a-c**). Along CD_go_, selectivity was larger, emerged faster, and persisted longer in the PT^lower^ population compared to PT^upper^ cells (**Fig. 6 b,c**). In the PT^lower^ population, significant selectivity emerged 24 ms following the go cue, faster than in the PT^upper^ population (46 ms) (**Fig. 6c**). The onset of the first detectable movement occurred approximately 50 ms after go cue onset (99% confidence interval = 38-56 ms). The coding dimension changed rapidly at the go cue in the PT^lower^ population, and more slowly, over several hundred milliseconds, in the PT^upper^ population (**EDFig. 10**). Individual PT^lower^ neurons displayed pronounced changes in selectivity at the go cue (**Fig. 6d**).

**Figure 6.**
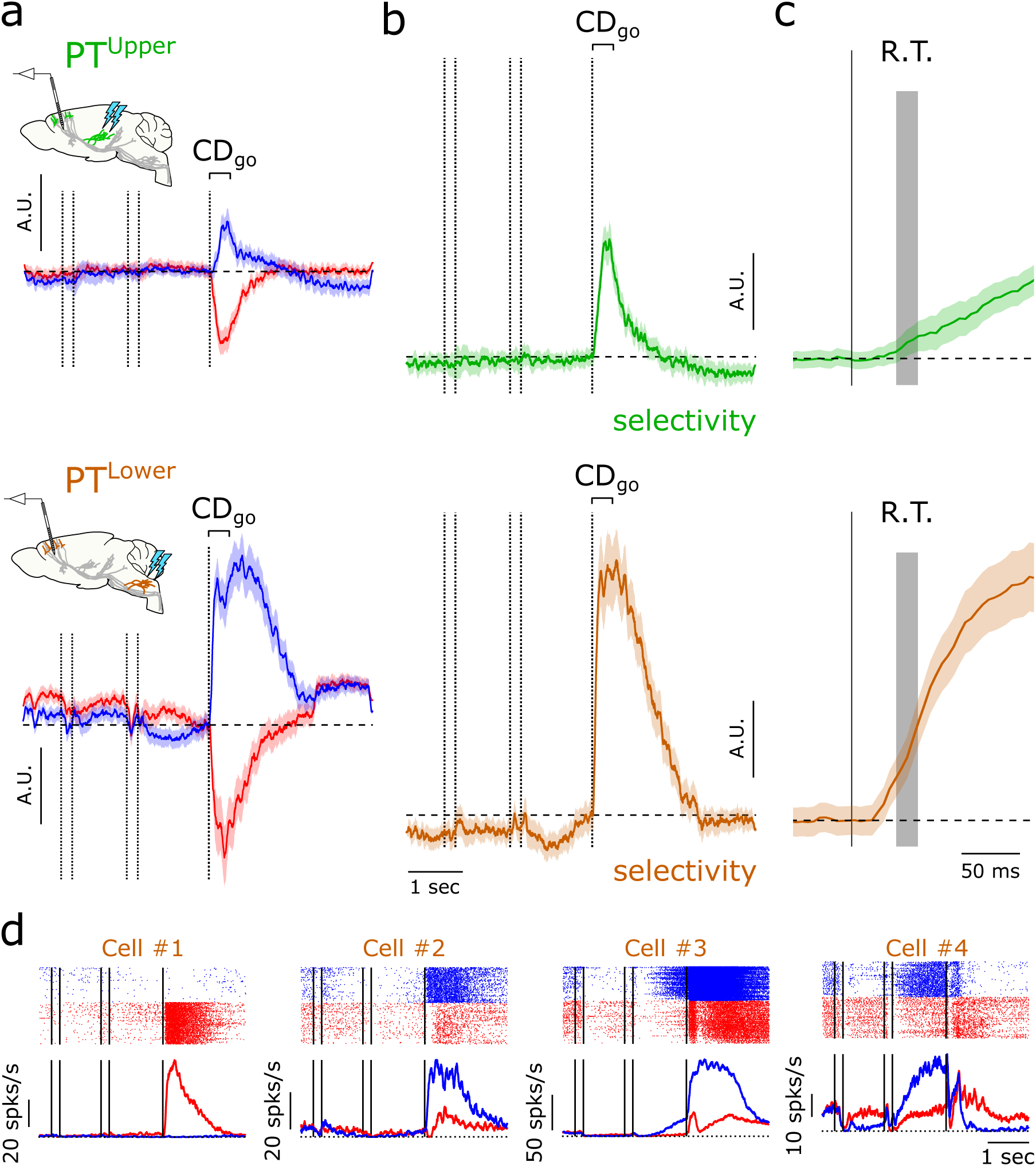
Movement commands in in PT^lower^ neurons. **a.** Time-course of the linear combination of neuronal activity that best differentiates trial types after the go cue (CD_go_) on lick right (blue) and lick left (red) trials for PT^upper^ (*top*) and PT^lower^ (*bottom*) neurons. **b.** Difference in CD_go_ projections on lick right and lick left trials (selectivity) in PT^upper^ (*top; green*) and PT^lower^ (*bottom; orange*) neurons.**c.** Data from (b) expanded around the go cue. Gray region indicates the distribution of session-averaged reaction times (earliest detected orofacial movement, *R.T.;* 99% confidence interval = 38-56 ms). Along the CD_go_, selectivity in PT^lower^ neurons emerged 24 ms following the go cue, faster than in the PT^upper^ population (46 ms) and consistent with a role in movement initiation. **d.** Example identified PT^lower^ neurons. *Top:* raster plots for correct lick right trials (blue) and lick left trials (red). *Bottom:* trial-averaged spike rates. Shaded regions in (a-c) represent 95% confidence intervals around the mean using hierarchical bootstrapping.

Each bout of licking consists of a sequence of directional tongue protrusions at a stereotyped frequency (approximately 8 Hz) (**Fig. 4,b**). Aligning PT^lower^ activity to the last lick in a bout revealed additional movement-related features. Selectivity along both the CD_late_ and the CD_go_ dimensions ceased with the offset of movement (**EDFig. 12a,b**), simultaneous with an abrupt change in the coding dimension (**EDFig. 12c**). This transition was not observed in the dynamics of the PT^upper^ population (**EDFig. 12a-c**). Indeed, examining the activity of single neurons in the PT^lower^ population revealed neurons that were strongly modulated at the go cue, at the offset of movement, or both (**EDFig. 12d**). These results show that subgroups of PT^lower^ neurons have activity patterns consistent with roles in initiating and/or terminating movements.

## DISCUSSION

Pyramidal tract (PT) neurons of motor cortex exhibit diverse activity patterns that are related to movement planning and execution^3,5,18,20,35^. We have shown that PT neurons in motor cortex comprise at least two cell types with distinct gene expression patterns, axonal projections, and specialized roles in motor control. PT^upper^ neurons connect with the thalamus and avoid motor centers in the medulla. PT^upper^ neuron activity represents a short-term memory that links sensory information and motor planning. PT^lower^ neurons avoid the thalamus and project to motor nuclei in the medulla. These neurons appear to control movement initiation and termination. PT neurons segregate into PT^upper^ and PT^lower^ types across the entirety of motor cortex (**EDFig. 4**). This organizing principle may extend to non-motor cortical areas and other mammals^36–38^.

Previous anatomical studies have suggested that collaterals of PT neurons projecting to motor centers also innervate the thalamus^14,39^, possibly providing an efference copy of motor commands for predicting the sensory and motor consequences of self-movement^15^. We show that neurons that project to motor centers do not project to the thalamus. Instead corticothalamic PT^upper^ neurons play more cognitive roles in motor preparation. The thalamus could still process efference copies of motor commands through an indirect, tectal pathway^40,41^. The thalamus also receives a projection from L6 corticothalamic neurons, but these neurons are sparsely active and uncoupled from PT neurons^16^. In addition, their weak synapses are thought to play a modulatory role in thalamic excitation^15^.

Cell type-specific recordings link representations of information with the neural circuit diagram, which is critical to understand the mechanisms of neural computation. The two PT neuron types express distinct behavior-related signals, implying that only a subset of the information represented in a cortical region is communicated to specific downstream structures^18,42,43^. Preparatory activity appeared early, and was persistent, in PT^upper^ neurons, whereas movement commands were observed in PT^lower^ neurons. At the same time, multiple signals were multiplexed within the population of PT^lower^ neurons. Preparatory activity emerged in this population during the delay epoch (along CD_late_) and persisted through the go cue and up to the termination of licking bouts. In the same cell type, and sometimes in the same individual cells (e.g. **Fig 6d**; *Cell #3),* activity was strongly modulated following the go cue along a different dimension (CD_go_), consistent with a movement command. A complete description of neural coding therefore requires measurements of neural populations of defined cell types.

In this study we restricted our analysis to ALM PT neurons and found two principal cell types. PT^upper^ neurons corresponded to two transcriptomic clusters (**Fig. 1**), separated by many differentially expressed genes (**Fig. 3a** and **EDFigs. 1,6**). Future studies linking detailed anatomy with transcriptional profiling might lead to further subdivision of these types. In addition to PT neurons, ALM harbors ten transcriptomic clusters corresponding to diverse IT neurons, which project to other cortical areas and the striatum^22^ (**Fig. 1**). A subset of these neurons connect the two ALM hemispheres via the corpus callosum, which is critical for robustness in preparatory activity^30^. Identifying their roles in movement will require experiments similar to those presented here, in addition to mapping the connections between these cell types.

## METHODS

### Animals

Mice used for scRNA-seq experiments in visual cortex and ALM are described in Ref [22]. Mice used for all other experiments are described in **EDTable 2** and **EDTable 3**. Mice were housed on a 12-hour light-dark cycle with ad libitum access to food and water, except during behavior (described in ***Mouse behavior***).

### Surgical procedures

All procedures were in accordance with protocols approved by the Janelia Research Campus Institutional Animal Care and Use Committee and Institutional Biosafety Committee. Mice were given buprenorphine HCl (0.1 mg/kg; Bedford Laboratories) and ketoprofen (5 mg/kg; Fort Dodge Animal Health) for post-operative analgesia and to reduce inflammation. Surgical procedures were carried out under 1-2% isoflurane anesthesia. Mice were placed in a stereotaxic headholder on a thermal blanket and their eyes covered with artificial tears (Rugby). Marcaine (0.05 mL, 0.5%) was injected under the skin covering the skull to be thinned. The periosteum was removed and the skull thinned overlying the sites of viral injection(s). For all injections, virus was injected using a manual volume displacement injector (MMO-220A, Narishige) connected to a glass pipette (5-000-2005, Drummond Scientific) pulled to a 30 μm tip (P-2000, Sutter Instruments) that was beveled to a sharp tip. Pipettes were back-filled with mineral oil and virus was front-loaded prior to injection. Pipettes were inserted through the thinned bone to the appropriate depth and virus injected at a rate of 10 nL/min. For electrophysiology, a fiber optic cannula (CFML12L05; Thorlabs) was implanted 200 μm above the virus injection target and a headbar implanted caudal to Bregma. Dental acrylic (Jet repair; Pearson Dental) was used to secure the optic fiber and headbar to the skull and protect exposed bone.

### Viral expression

All viruses used in these experiments were adeno-associated virus (AAV) produced in the Janelia Research Campus (JRC) Virus Shared Resource. Viruses used for scRNA-Seq experiments are described in Ref [22]. For all other experiments, viruses incorporated the AAV2-retro capsid^28^, with the exception of axonal reconstruction experiments (**Fig. 2a,b** and **EDFig. 2**), which used AAV2 serotype 1. Viruses used, viral titers, injection volumes, injection coordinates, and associated experiments are described in **EDTable 2** and **EDTable 4**.

### scRNA-Seq

Single-cell RNA-Seq data (**Fig. 1**, **Fig. 2e,f**, **EDFig. 1**, and **EFig. 5**), was collected and analyzed according to Ref [22]. A total of 9,035 scRNA-Seq transcriptomes were measured from ALM neurons and 12,714 from V1 neurons. To collect individual cells, we used layer-enriching dissections from brains of *pan*-neuronal, *pan*-excitatory or *pan*-inhibitory recombinase lines crossed to recombinase reporters (4,506 cells). This dataset was supplemented by 3,196 cells isolated from other recombinase lines. Dissections without layer enrichment, or multiple layers combined, were employed for lines with sparse labeling. Additional recombinase lines were selected to capture cellular diversity that was suggested by ongoing analysis of the data. 1,333 additional cells were derived from viral retrograde labeling, with the goal to establish correspondence between transcriptomic types and projection properties. PT neurons were harvested from retrogradely-labeled brains (n=63 medulla injection; n=94 thalamus; n=100 pons; n=92 superior colliculus; n=2 amygdala; n=2 zona incerta; 5 retrosplenial cortex) and recombinase crosses (n=10). In all cases, L5 was micro-dissected to isolate PT neurons from, for example, thalamus-labeled L6 CT neurons.

### scRNA-Seq analysis

Transcriptomic features, extracted by weighted gene co-expression network analysis^23^, were clustered in an iterative and bootstrapped manner. The output of this procedure is a co-clustering matrix, which shows the frequency with which any cell clusters with any other cell in 100 bootstrapped iterative clustering rounds. The transcriptomic clusters (i.e. putative cell types) are defined by ‘cutting’ the co-clustering matrix to derive membership of each cell to a cluster. Each cell’s membership is tested post-clustering by classification algorithms to assign core vs. intermediate identity to cells ^44^. Cells that are reliably assigned to only one cluster are called ‘core’ cells 19,195), cells that get assigned to more than one cluster (typically two), are ‘intermediate’ cells 2,554 cells). Each cluster was named based on known or newly discovered differentially expressed markers. ALM scRNA-Seq transcriptomes clustered into 21 glutamatergic, 49 GABAergic and 14 non-neuronal types. Cells labeled retrogradely from thalamus, medulla, superior colliculus or pons were submitted to scRNA-Seq, with the results mapping onto the pre-established taxonomy.

### Single-cell axonal reconstruction

The axons of single neurons were labeled and imaged as described (**Fig. 2a,b** and **EDFig. 2**)^27^. Neurons in motor cortex were sparsely labeled using a viral vector encoding either eGFP or tdTomato. At least 3 weeks following virus injection, mice were perfused and brains extracted. Brains were embedded in gelatin and cleared using a combination of DMSO and D-sorbitol. The full volume of each brain was imaged at submicron resolution using an automated block-face two-photon microscope with integrated vibratome. Full-brain datasets were approximately 20-30 TB in size and were stitched and rendered using a custom bioinformatics pipeline^27^. Each individual neuron was reconstructed manually in three-dimensions using the Janelia Workstation^45^ by two independent annotators who were blinded to all analyses. Consensus reconstructions were determined by resolving discrepancies (generally <5%) between annotators. Each dataset was registered to the Allen Common Coordinate Framework using landmark based registration (3DSlicer, Landmark Registration module) and a thin plate spline warp determined between the two image spaces. Neuronal reconstructions were then projected into the reference space to determine the brain area associated with each axonal segment, branch point, and terminus.

### Histology and imaging

At least three weeks following viral injections, mice were trans-cardially perfused with PBS (>20 mL) followed by 4% paraformaldehyde (>20 mL). Brains were post-fixed overnight. For immunolabeling (**Fig. 2c,d** and **EDFig. 4**), brains were transferred to a 20% sucrose solution for cryoprotection and sectioned coronally at 50 μm on a freezing microtome. In all other cases (**EDFig. 3**), brains were sectioned at 50 or 100 μm on a vibratome (VT1200; Leica Biosystems). Sections were processed using standard immunohistochemical techniques and imaged as described previously (**Fig. 2c,d** and **EDFigs. 3,4**)^46^. Brightness and contrast were adjusted manually to match approximate luminance values across imaging experiments in a linear fashion using ImageJ/Fiji.

### Bulk RNA-Seq

Cells back-labeled from thalamus, medulla, pons and superior colliculus (SC) with GFP or tdTomato were collected by manual cell sorting, as described (**EDFig. 5** and **EDFig. 6**)^47^. 50-70 cells were collected per sample, with 7 experimental replicates performed for each. Cells were isolated by manual cell sorting on a fluorescence dissecting scope, following micro-dissection, trituration and enzymatic digestion^48^. Following pooling and lysis, total RNA was extracted by Picopure kit (KIT0204; Thermo-Fisher). Amplified DNA was produced using Ovation RNA-Seq System V2 kit (#7102; NuGEN), fragmented to ∼200bp, ligated to Illumina sequencing adaptors with Encore Rapid kit (#0314; NuGEN), and sequenced on an Illumina HiSeq 2500 with 4-fold multiplexing (single end, 100bp read length).

### Bulk RNA-Seq analysis

Adaptor sequences (AGATCGGAAGAGCACACGTCTGAACTCCAGTCAC) were removed from reads using Trimmomatic 0.36^49^, mapped using STAR 2.5.3a^50^ to the Ensembl mouse genome GRCm38.p5, release 90 (https://www.ensembl.org\). Mapped reads were normalized to the total number of reads per sample (counts per million). Differential-expression (DE) criteria were: false-discovery-rate < 5%; log2-fold-change > 2.0; sample CPM mean > 10.0 in 3 or more replicates. Principal components analysis of the 7 thalamus-projecting and 7 medulla-projecting replicates showed that a single replicate from each behaved as an outlier. Thus, for further analysis, 6 replicates were considered for each. For selection of potential marker genes, non-coding RNAs and gene models were removed from the DE gene lists. For visualization, reads per kilobase of transcript per million mapped reads (RPKM) were used.

Similar numbers of genes were detected in each of the thalamus and medulla bulk RNA-Seq samples (replicate-averaged #detected genes: 12,317 and 12,363 respectively). These numbers were slightly higher than the genes detected from scRNA-Seq (thalamus: 11,435; medulla: 11,573; pons: 9,486; and SC: 9,742, respectively).

### *In situ* hybridization

Animals were perfused with 4% PFA and brains were post fixed in 4% PFA overnight at 4 degrees C. Brains were then rinsed in PBS and cryoprotected in 30% sucrose, and 20 μm thick sections were cut on a cryostat. smFISH followed by IHC was performed on fixed frozen tissue from mice injected with AAV2-retro-mRuby2-smFP-FLAG either in thalamus or in medulla (**EDTables 2–4**) as per protocols for fixed frozen tissue using proprietary probes from Advanced Cell Diagnostics (ACDBio). Probes used in this study (Mm-Npnt: Cat# 316771; Mm-Slco2a1: Cat# 485041) were detected using propriety detection reagent (RNAscope Fluorescent Multiplex Detection Reagents Cat # 320851), using Amp4 Alt B (Atto 550). Following smFISH, sections were rinsed in PBS, and blocking buffer (2% BSA and 0.3% serum) was applied for 5 minutes. Primary antibody (Sigma-Millipore F1804) was diluted in the blocking buffer (1:100) and incubated overnight at 4 degrees C. Sections were rinsed in PBS 3 times (5 minutes) and incubated with secondary antibody Goat anti-mouse AF 488 (A-11001, Invitrogen; diluted 1:100 in blocking buffer) at room temperature for 2 hrs. Sections were then rinsed in PBS and cover slipped with Vectashield containing DAPI (H-1500; VectorLabs). Images for display were acquired as a single plane on a Zeiss 880 inverted confocal microscope, using a 63X/1.4NA objective. Images for quantification were acquired on 7μm thick stacks using a 40X oil immersion objective/1.3NA (pixel size, 0.11 × 0.11 μm). Laser power was adjusted across sections to achieve maximum dynamic range. Punctate mRNA signal was quantified on cell volumes from maximum intensity projection of the Z-stacks. Brightness and contrast were adjusted manually to match approximate luminance values across imaging experiments in a linear fashion using ImageJ/Fiji.

### Mouse behavior

Mice were water restricted and housed on a 12-hour reverse light-dark cycle with testing during the dark phase. On days in which mice were not trained, they received 1 mL of water. Behavioral experiments lasted one to two hours per day, during which period they consumed their daily water intake (∼ 0.5 to 1.0 mL). Mice unable to sustain stable body weight were given supplementary water. Mice were trained using operant conditioning as previously described^26,51^ until reaching behavioral criterion (>75% trials correct). At the beginning of each trial, a vertical pole moved into place adjacent to the whisker pad and in reach of the whiskers (200 ms travel time). The pole remained in this position for 1.0 sec and then was retracted (200 ms travel time). The sample epoch was defined as the 1.0 sec during which the pole was in range of the whiskers and stationary. After the pole was removed, the mouse was trained to refrain from licking for an additional 1.3 sec (delay epoch) before an auditory ‘go cue’ (pure tone, 3.4 kHz, 0.1 s duration) instructed the mouse to lick (reward epoch). Premature licks during the sample or delay epoch resulted in a restart of the requisite epoch and these trials were excluded from all analyses. Licking the correct lickport after the go cue led to a small water reward (3 μL). Licking the incorrect lickport triggered a timeout (2-10 sec). Trials in which mice did not lick within a 1.5 sec window after the go cue were rare and typically occurred at the end of a session.

### Videography

High-speed video was acquired at 400 Hz from below and to the side of the mouse using CCD cameras (CM3-U3-13Y3M; FLIR) with a 4-12mm focal length lens (12VM412ASIR; Tamron). Camera data were acquired using BIAS (IORodeo). Reaction times were determined by measuring the luminance change in a small ROI manually placed just below the jaw in side-view movies. Luminance traces were averaged across all of the trials within each session (453 ± 79 mean ± s.d.; range: 295 - 551). 95% confidence intervals for reaction time were calculated by bootstrapping session means across mice (n = 3) and sessions (n = 14). Little inter-animal variation was observed in session-averaged reaction time.

### Electrophysiology

A small craniotomy (diameter, 0.5 - 1 mm) was made over ALM one day prior to the first recording session. Extracellular spikes were recorded using silicon probes containing two shanks each with 32 channels with 25 μm spacing (H2; Cambridge Neurotech). The 64 channel voltage signals were multiplexed, recorded on a PCI6133 board (National instrument), and digitized at 14 bits. The signals were demultiplexed into the 32 voltage traces, sampled at 25 kHz and stored for offline analyses. 4-7 recordings were made from each craniotomy on consecutive days. Recording depth was inferred from manipulator readings without compensation for cortical curvature. The tissue was allowed to settle for 10 minutes prior to recording.

To optogenetically tag PT^upper^ and PT^lower^ neurons during recording, we expressed ChR2(H134R)-YFP selectively in each population using a viral injection into the medulla or thalamus as described above. During each recording session, >1200 optical stimuli were delivered at 4 Hz just prior to and following the behavioral session. Stimuli were 0.1-0.5 ms at 80-100 mW (measured just before the implanted fiber optic cannula). Reliable antidromic activation was observed in 1-6 units per session. Due to the proximity of the cerebral peduncle, subthalamic nucleus, and zona incerta, which contain projections from both cell types, a smaller viral injection was performed in the thalamus (50 nL) than in the medulla (50-200 nL), resulting in fewer tagged PT^upper^ neurons and reduced throughput. Across all sessions recording PT^upper^ neurons, mouse performance was 85.4 ± 7.8% on lick left trials and 89.3 ± 9.4% on lick right trials (mean ± s.d.; n = 37 sessions in 8 mice; left hemisphere). In PT^lower^ neuron recordings, performance was 83.5 ± 6.3% on lick left trials and 86.3 ± 8.2% on lick right trials (n = 23 sessions in 4 mice; both hemispheres). Mice performed a median of 107 and 105 correct trials on lick left and lick right trials respectively in PT^upper^ recordings and 117 and 108 correct trials during PT^lower^ recordings. PT neurons recorded from the left and right ALM did not differ qualitatively and were combined in all further analysis to increase statistical power.

### Electrophysiology data analysis

Extracellular recorded traces were band-pass filtered (300 – 4500 Hz; 2^nd^ order Butterworth filter) and the common mode on ±4 sites was subtracted from each channel. Events were detected using JRClust^52^ and spikes from tagged units (n = 143) were sorted manually using a custom program written in MATLAB. Extreme care was taken to restrict analysis to units that could be sorted with a low number of false positive spikes (mean ISIs less than 2 ms = 0.02%; **EDFig. 7**) so that neuronal responses could be faithfully attributed to the correct cell type. Despite this, spike rates were similar to that measured in other studies recording extracellular from ALM in the same task ^18^ indicating that the false negative spike rate remained low. Information from 4-7 adjacent used for sorting each individual unit. Units recorded during behavioral sessions in which performance was not greater than 65% for both trial types were excluded from the dataset. Additionally, units recorded during behavioral sessions with less than 50 correct trials of each type were excluded. 13 units were rejected based on these criteria and 130 were kept for further analysis. Unit depths (**Fig. 4e**) were inferred from manipulator readings only without correction for the angle between the electrode penetration and the orientation of cortical layers. Collision tests were performed for all tagged units to confirm axonal projections to the thalamus or medulla^18,53^ (**Fig. 4d**). Trial-averaged spike rates were calculated in 5 ms time bins and filtered using a causal 50 ms boxcar filter.

Coding dimension vectors (**Figs. 5,6** and **EDFigs. 9,12**), ***CD,*** were calculated according to equations 1 and 2.

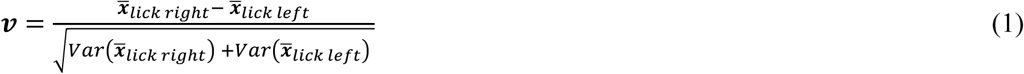

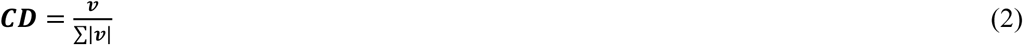

For each unit, the mean difference in spike rate between lick right, *x̅_lick right_*, and lick left trials, *x̅_lick left_*, was calculated across a 400 ms time interval. This vector was divided by the square root of the sum of the across-trial variances of spike rate for each trial type. The resulting vector (***ν***) was then normalized by its L1 (taxicab) norm so that projections would not scale with vector length (# cells in population) to produce the coding dimension, ***CD***. Coding dimensions were calculated in the first 400 ms of the sample epoch (CD_early_; 2.5 to 2.1 seconds before the go cue) the last 400 ms of the delay epoch (CD_late_; 0.4 to 0.0 seconds before the go cue) and the first 400 ms of the response epoch (CD_go_; 0.0 to 0.4 seconds after the go cue). CD_go_ was further orthogonalized to CD_late_ using the Gram-Schmidt process to remove the component of CD_late_ that persisted through the response epoch from CD_go_. In all cases, coding dimensions were calculated separately for the PT^upper^ and PT^lower^ populations.

Projections of the activity of each PT population along the coding dimensions (***p***_*lick right*_, ***p***_*lick left*_) were obtained according to equations 3 and 4:

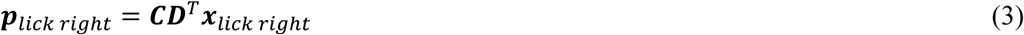

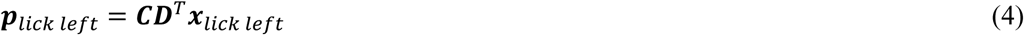

Selectivity, ***S***, along each dimension was calculated as:

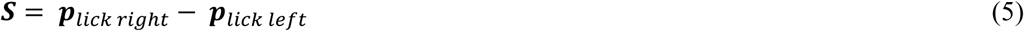

In all cases (**Figs. 5,6** and **EDFigs. 9,12**), the illustrated projections are the results of a hierarchical bootstrapping procedure. For each population, 100 projections were calculated, each using N randomly chosen neurons (with replacement), where N was the number of neurons in the population. For each neuron, 50 correct trials of each trial type were randomly selected with replacement. In all figures, solid lines and shaded areas represent the median and standard deviation of all repetitions.

To create matrices of the coding dimension correlation across time (**EDFig. 10**), coding dimensions were calculated at each time point from the trial-averaged spike rates of all neurons within a population (5 ms time bins, filtered using a causal 50 ms boxcar filter), as above, but were normalized by their Euclidean norms to produce unit vectors. Correlation matrices represent the inner product of coding dimension vectors at each pair of time points.

Single-cell trial type decoding accuracy (**EDFig. 11**) was determined using the average spike rate during the first 400 ms of the sample epoch (2.5 to 2.1 seconds before the go cue). A spike rate threshold was determined that best distinguished lick right trials from lick left trials (maximal accuracy) with accuracy defined as the proportion of trials correctly classified. Accuracy was >=50% by definition. Shaded regions in around the cumulative distribution function in **EDFig. 11b,e** represent standard error, estimated using Greenwood’s formula. In **EDFig. 11c,f**, confidence intervals represent standard errors (bootstrap) and the gray shaded region represents ±1 standard error of the expected value after shuffling cell type labels (100,000 repetitions).

### Code availability

All analysis code used in this study is available upon request.

## ACKNOWLEDGEMENTS

We thank Andy Lemire, Kshama Aswath for single-cell sorting and bulk RNA-Seq, Sara Lindo for stereotaxic surgeries, Damian Kao for help with bulk RNA-Seq analysis, and Nuo Li and Hidehiko Inagaki for help with electrophysiological recordings and helpful discussion. We thank Mark Cembrowski, Erik Bloss and Frederick Henry for helpful discussion. Hidehiko Inagaki, Murray Sherman, Sandro Romani, Liqun Luo, Gordon Shepherd, and Tim Wang provided comments on the manuscript.

## AUTHOR CONTRIBUTIONS

MNE, KS, SV, LL, BT, and HZ; conception and design of the experiments. BT, TNN, LTG and HZ; scRNA-seq experiments, with material provided by MNE and SV. MNE, EB, JW, and JC; single neuron reconstructions. SV; in situ experiments and analysis. MNE; electrophysiology experiments. MNE, SV, and CRG; anatomical experiments. MNE, SV, VM, LG, TN, KS and LL; data analysis. MNE, KS, and LL; wrote the paper with input from all authors.

## DATA AVAILABILITY

Single cell transcriptomic data will be available through GEO. Bulk RNA-Seq data will be deposited in the NCBI Sequence Read Archive. Electrophysiology data sets will be shared at CRCNS.ORG in the NWB format.

## EXTENDED DATA FIGURE LEGENDS

**Extended data figure 1.**
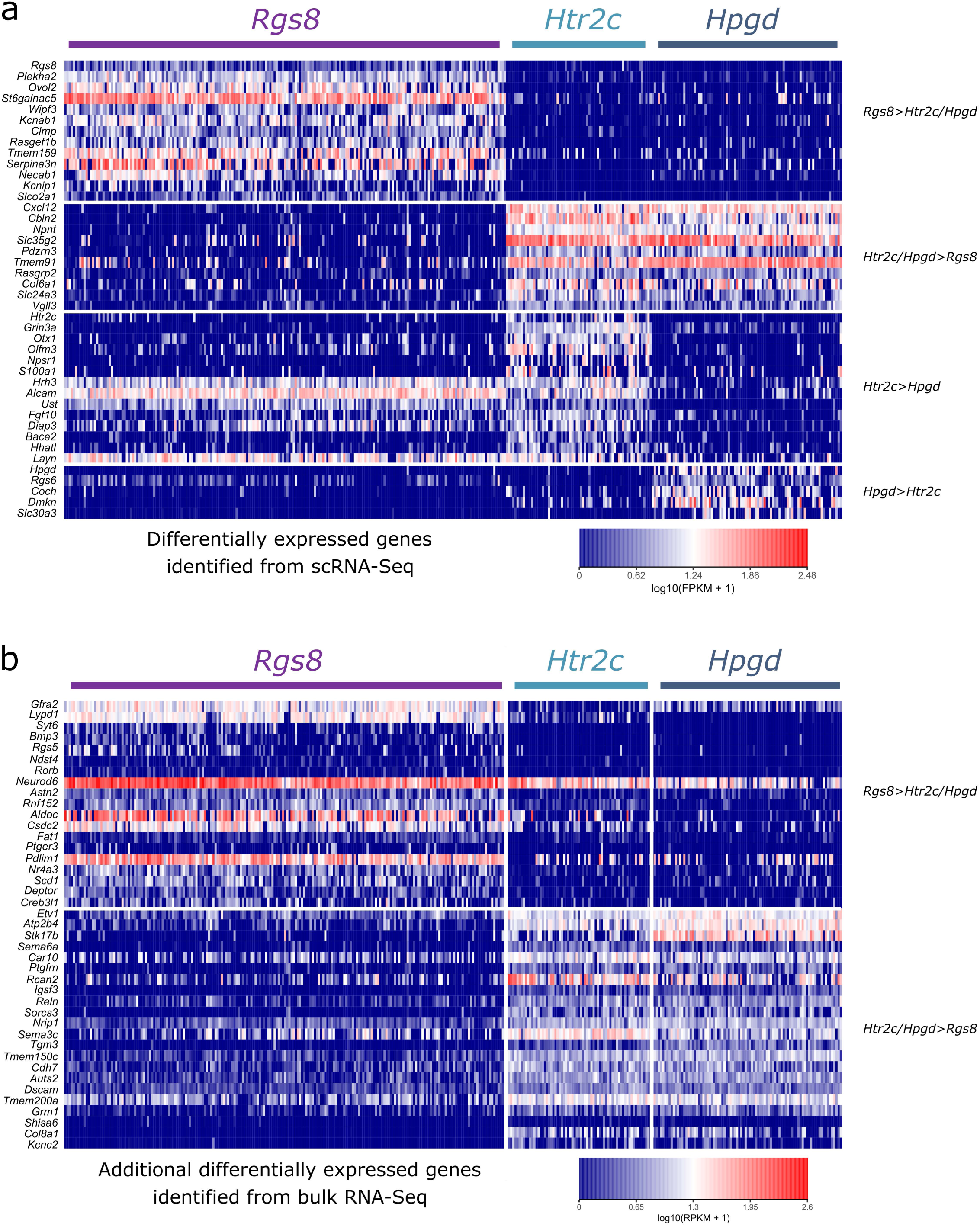
Differentially expressed genes in single cell RNA-seq PT neuron expression. **a.** Heat map of expression of differentially expressed genes. Columns in the heat map represent individual cells, grouped by cluster (Rgs8, *n*=209; Htr2c, *n*=69; Hpgd, *n*=90). Rows represent genes selected by the clustering algorithm to represent transcriptomic branch points between clusters. Color shows transcript expression in fragments per kilobase of transcript per million mapped reads (FPKM), shown on a log-scale. Pure blue shows cells with 0 transcripts of a given gene; pure red shows cells with maximal expression (approximately 300 FPKM). **b.** Expression data for the same cells in (a) for genes identified from bulk RNA-Seq as differentially expressed between the *Rgs8* and *Hpgd/Htr2c* clusters.

**Extended data figure 2.**
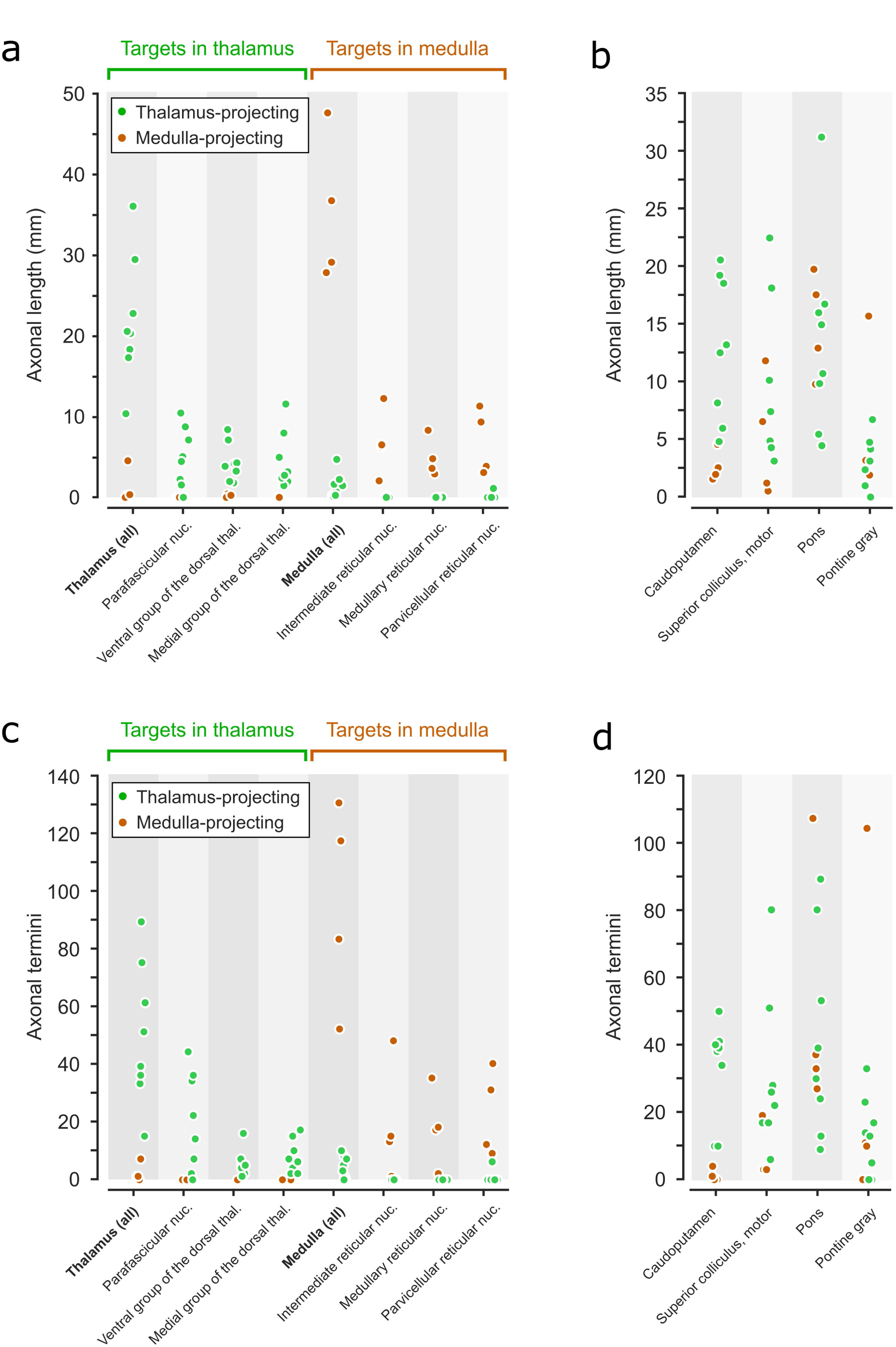
Distribution of axonal projections. **a.** Axonal length within thalamic and medullary targets for thalamus-targeting PT neurons (n = 8; *green*) and medulla-targeting PT neurons (n = 4; *orange*). **b.** Axonal lengths within other selected PT targets. **c.** Axonal termini within thalamic and medullary targets. **d.** Axonal termini within other selected PT targets.

**Extended data figure 3.**
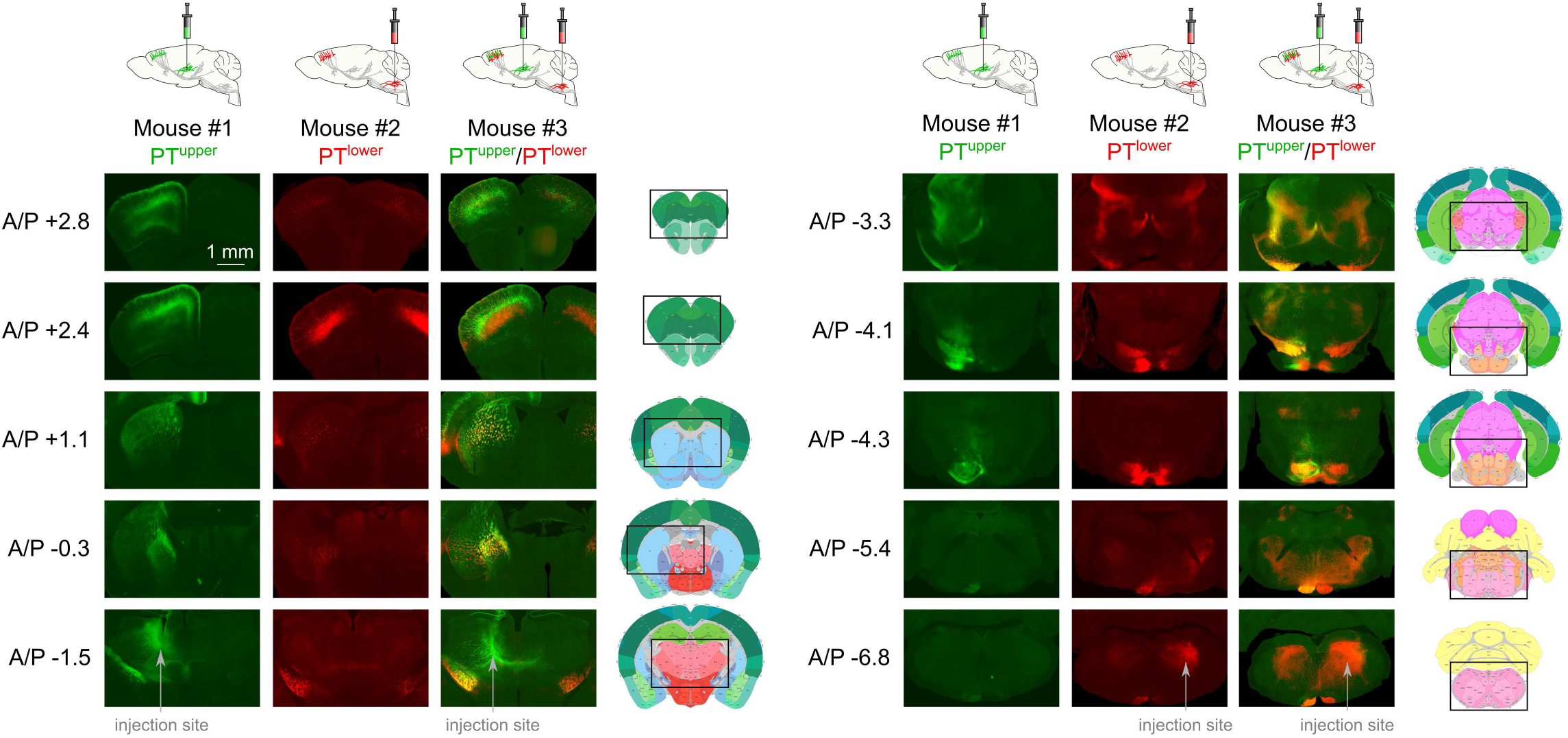
Anterograde anatomy: targets of PT^upper^ and PT^lower^ populations. Groups of cortical neurons were labeled from the thalamus (PT^upper^; Mouse #1), the medulla (PT^lower^; Mouse #2), or both targets (Mouse #3) using AAVretro expressing spectrally-distinct fluorescent proteins. Top, schematics of the labeling procedures. Left, rostro-caudal level (relative to Bregma). Right, imaged area is indicated on annotated coronal sections taken from the Allen Reference Atlas at the corresponding rostro-caudal level. Both types extended axon collaterals to motor-related SC, but to different parts. Axons from PT^upper^ cells were apparent throughout all SC layers, with a particularly dense projection to the ventrolateral aspect, whereas PT^lower^ neurons were restricted to the ventral superior colliculus and were concentrated more caudally. Both groups projected to the pons, particularly the pontine gray, but with terminations in largely non-overlapping zones. PT^upper^ cells exclusively projected to the globus pallidus external segment and broadly targeted the dorsal, lateral and ventral striatum. PT^lower^ cells projected sparsely to the lateral striatum. PT^lower^ neurons projected to the central amygdala and parasubthalamic nucleus, although these projections arise from cortical neurons outside of ALM (not shown). PT^lower^ neurons also made up the majority of the projection to the red nucleus, parabrachial nucleus, substantia nigra pars compacta, motor and sensory trigeminal nuclei in the hindbrain, and the pyramidal tract. Both cell types extended axon collaterals locally within the same sublamina as their somata, the subthalamic nucleus, zona incerta, and the midbrain reticular nucleus. PT^upper^ cells appeared to project more broadly to layer 1 in motor cortex. Mouse #1 and #2 were used for electrophysiological recordings and in both cases projections are labeled with ChR2.

**Extended data figure 4.**
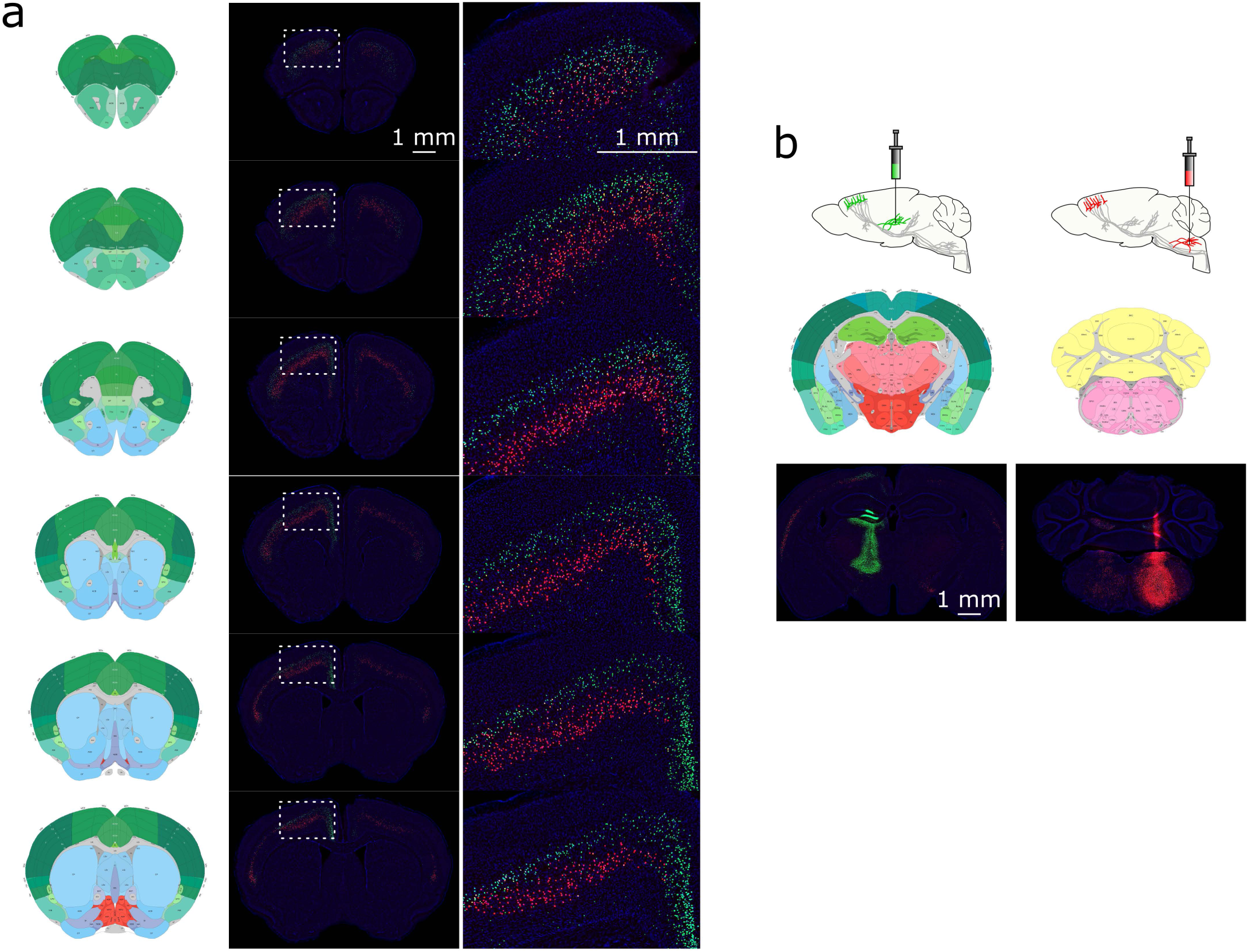
Distribution of thalamus- and medulla-projecting PT neurons. **a.** The nuclei of PT neurons were labeled from the thalamus (green cells) and medulla (red cells) using AAVretro. Thalamus-projecting PT neurons are in upper L5b throughout motor cortex, whereas medulla-projecting PT neurons are in deep L5b. Schematics to the left of each image set are annotated coronal sections (Allen Reference Atlas) at approximately the same rostro-caudal level. **b.** AA Vretro injection sites in the thalamus (*left*) and medulla (*right*).

**Extended data figure 5.**
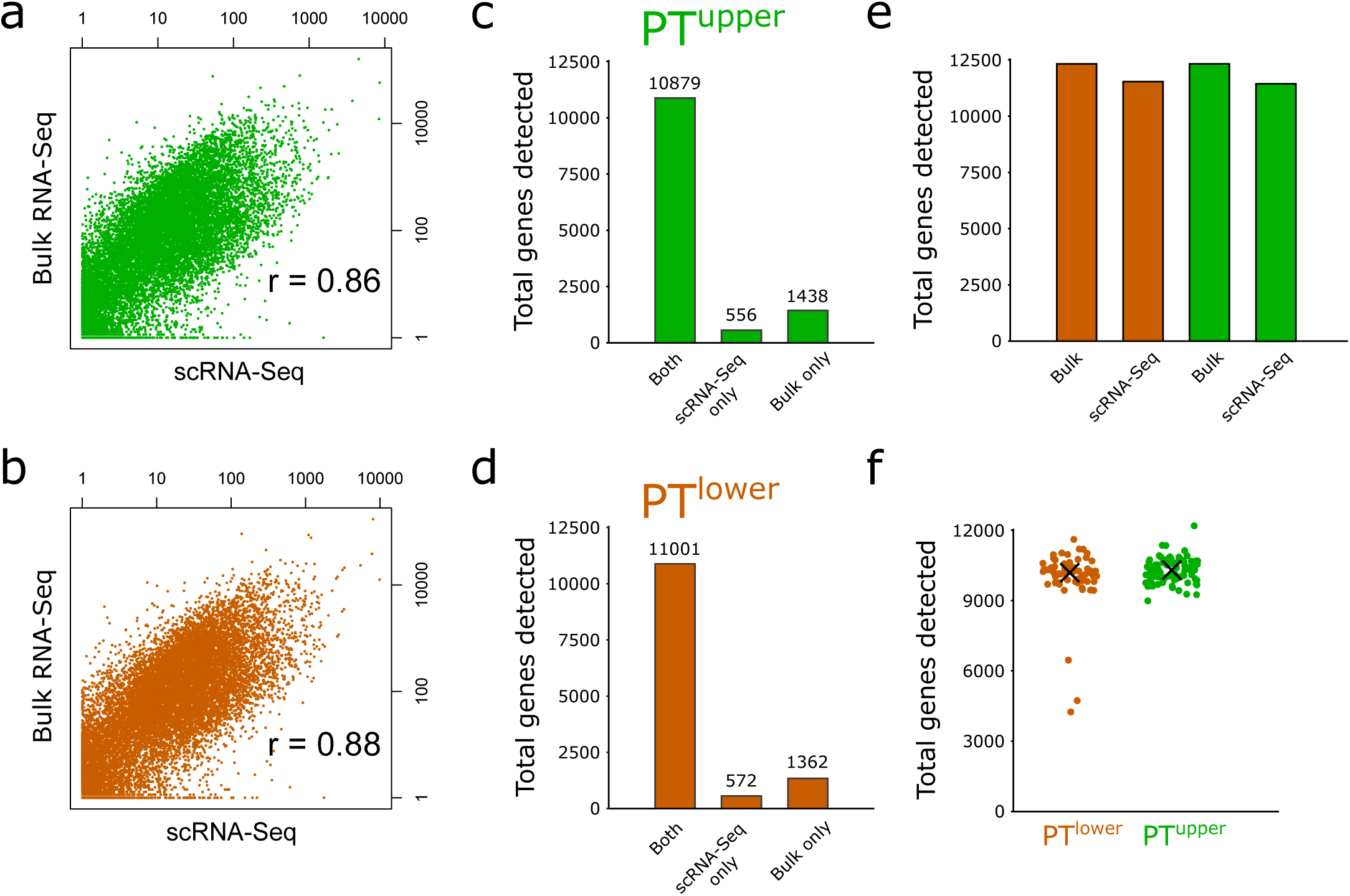
Comparison of single cell RNA-Seq and bulk RNA-Seq. **a.** Scatter plot of PT^upper^ gene expression (measured as FPKM + 1, shown in log-scale) from scRNA-Seq (x-axis) and bulk RNA-Seq (y-axis) (r = 0.86). **b.** Same as (a) for PT^lower^. **c.** Number of genes detected (Methods), from PT^upper^ neurons in bulk RNA-Seq and the union of scRNA-Seq measurements. The majority of genes were detected by both methods. **d.** Same as (c) for PT^lower^. **e.** Total genes detected by bulk RNA-Seq and scRNA-Seq in PT^upper^ and PT^lower^ neurons. **f.** Number of genes detected in scRNA-Seq for each PT^upper^ and PT^lower^ neuron (‘X’, median; PT^upper^, 9936 genes; PT^lower^, 9865 genes).

**Extended data figure 6.**
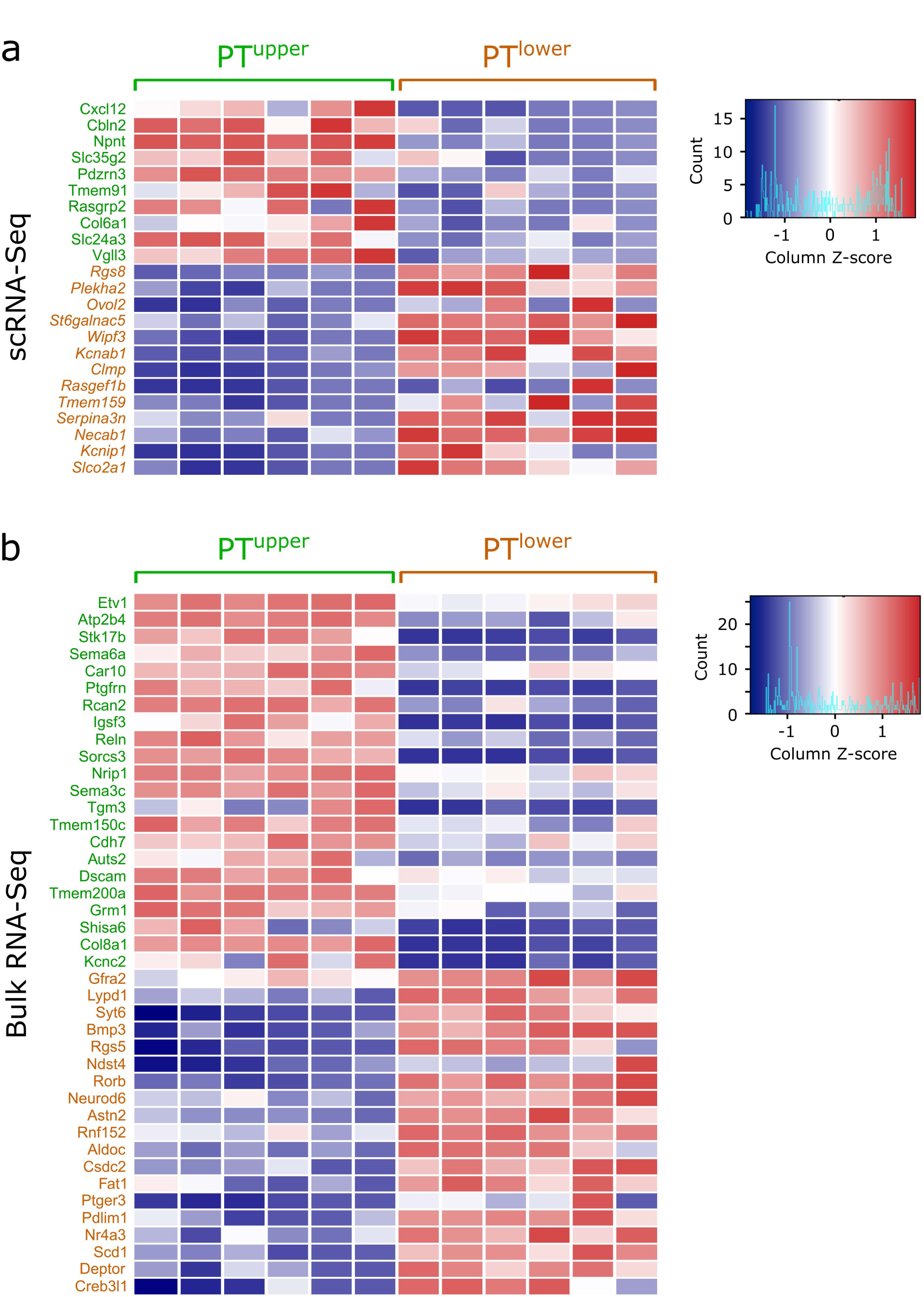
Differentially expressed (DE) genes in PT^upper^ and PT^lower^ cells, based on bulk RNA-seq. **a.** Genes identified as DE in scRNA-Seq, assessed in the bulk RNA-Seq data. Rows represent genes, colored by differential scRNA-Seq expression in PT^upper^ (green) or PT^lower^ (brown). Columns in the heat map represent transcript expression in individual replicates, 6 for each of PT^upper^ and PT^lower^. Colors show transcript intensity, as reflected from reads per kilobase of transcript per million mapped reads (RPKM), shown as Z-score. Pure blue shows replicates with very low expression (Z-score = −2); pure red shows replicates with very high expression (Z-score = +2). Bottom, distribution of z-scores. **b.** Same plot, for genes identified as DE from bulk RNA-Seq.

**Extended data figure 7.**
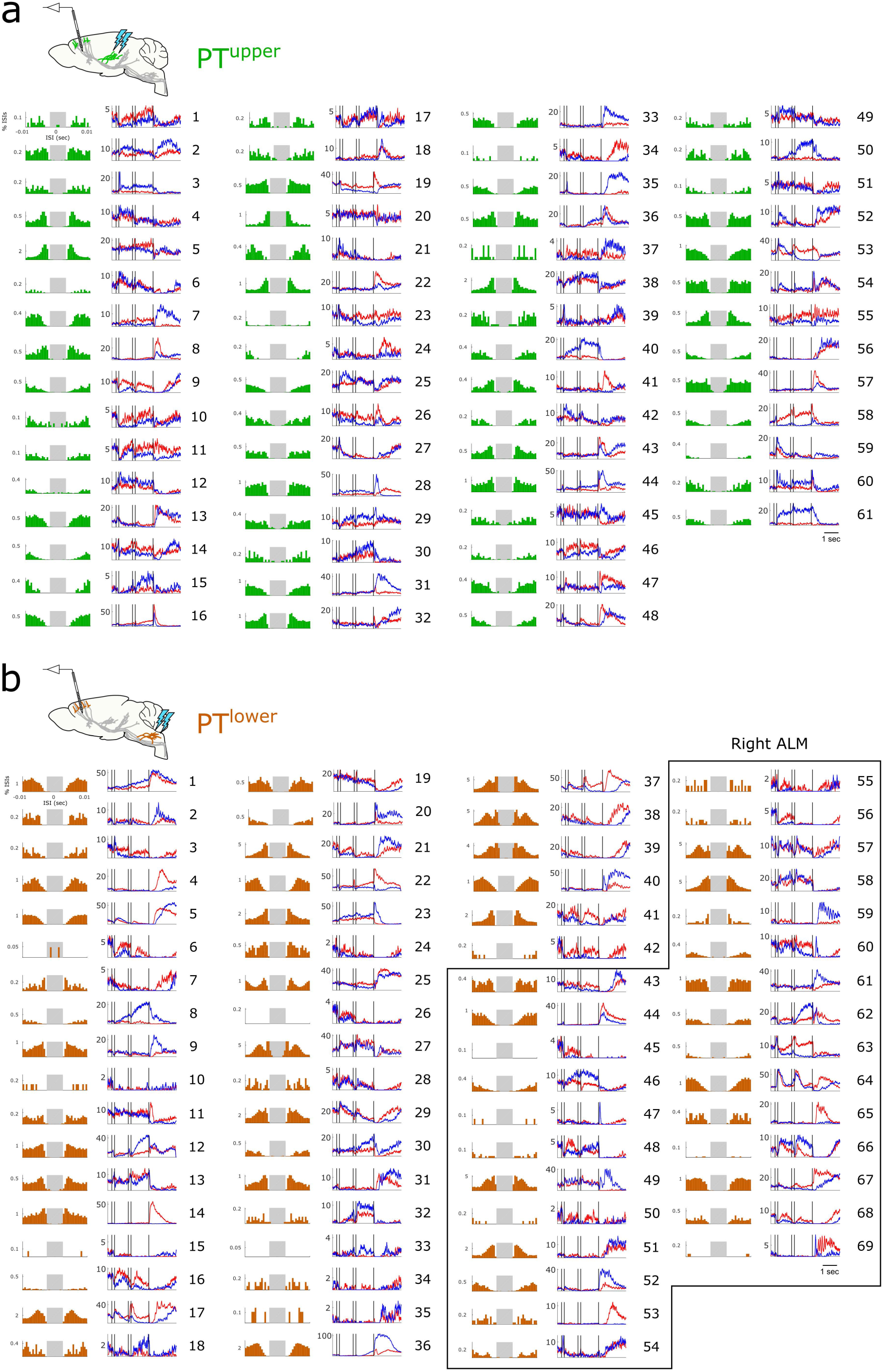
Electrophysiology and trial-averaged spike rates for all identified PT neurons. **a.** PT^upper^ neurons. Left, inter-spike interval (ISI) histograms; right, trial-averaged activity on lick right (*blue*) and lick left trials (*red*). Gray shaded area in ISI histograms represents the interval of -2.5 ms to 2.5 ms. **b.** PT^lower^ neurons. Boxed region indicates neurons recorded in the right ALM (ipsilateral to injection site in medulla). All other neurons were recorded in the left ALM (contralateral to injection site).

**Extended data figure 8.**
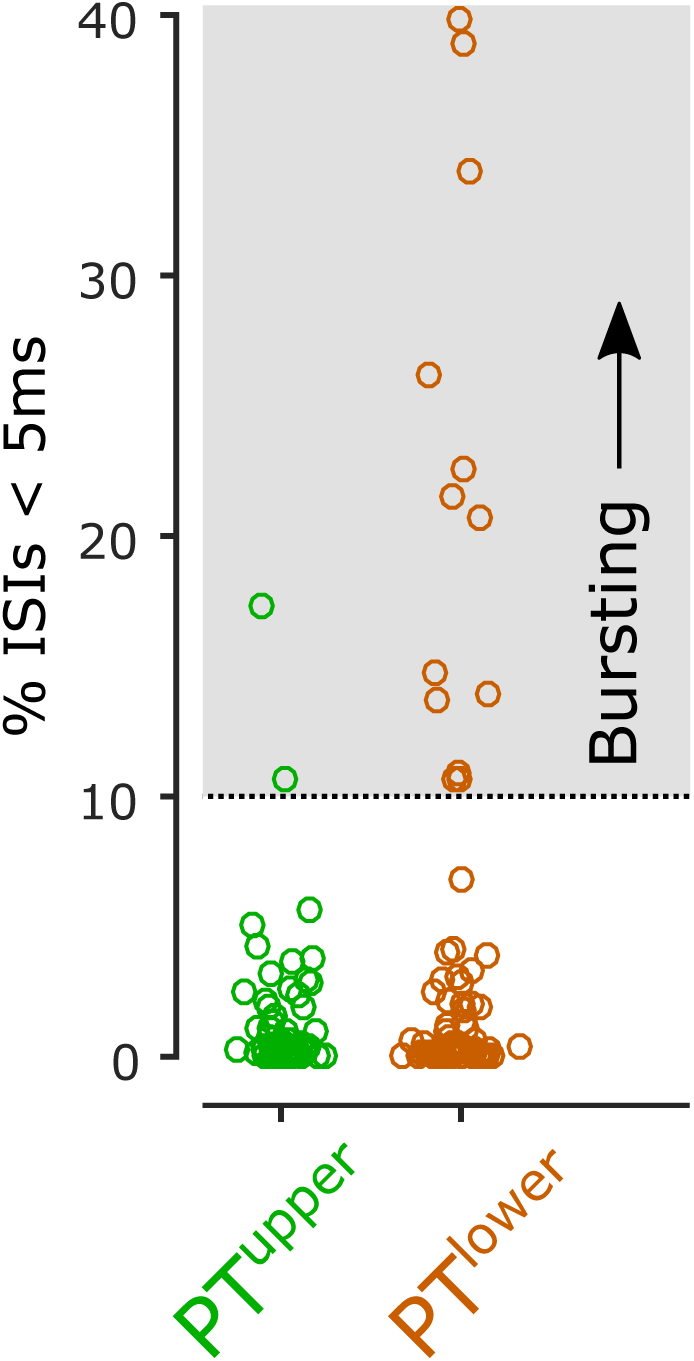
Bursting was detected predominantly in PT^lower^ neurons. Bursting cells (cells in which greater than 10% of inter-spike intervals were less than 5 ms) were rare in the PT^upper^ population (3.3%) and more common in the PT^lower^ population (18.8%; p = 0.006).

**Extended data figure 9.**
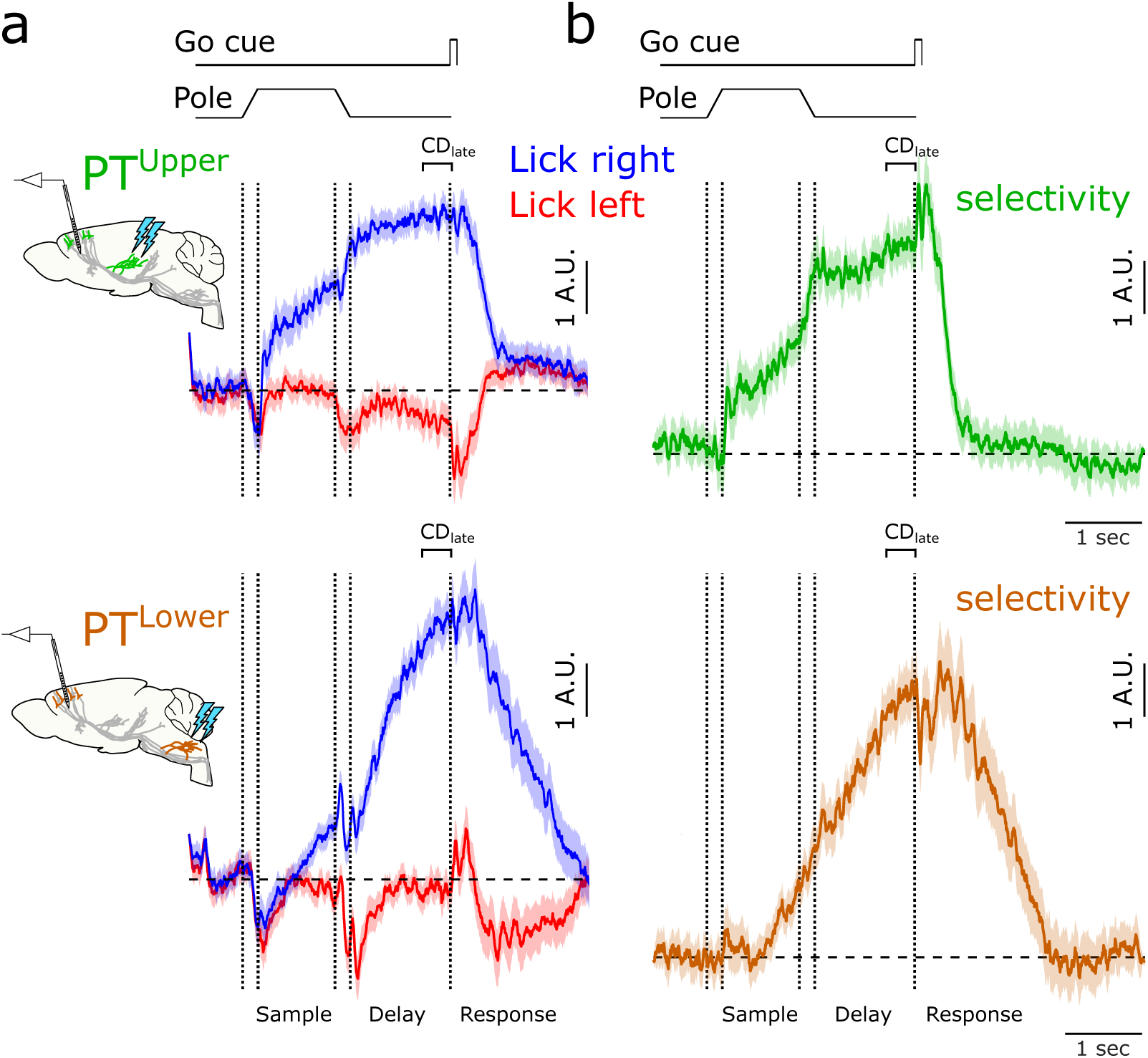
Preparatory activity in the late delay epoch. **a.** Time course of the linear combination of neuronal activity that best differentiates trial types in the 400 ms immediately prior to the go cue (late coding dimension; CD_late_) on lick right (*blue*) and lick left (*red*) trials for PT^upper^ (*top*) and PT^lower^ (*bottom*) neurons. **b.** Difference in CD_late_ projections on lick right and lick left trials (selectivity) in each population. Selectivity along CD_late_ is present in both populations, and persists after the go cue, but does is not strongly modulated during movement initiation.

**Extended data figure 10.**
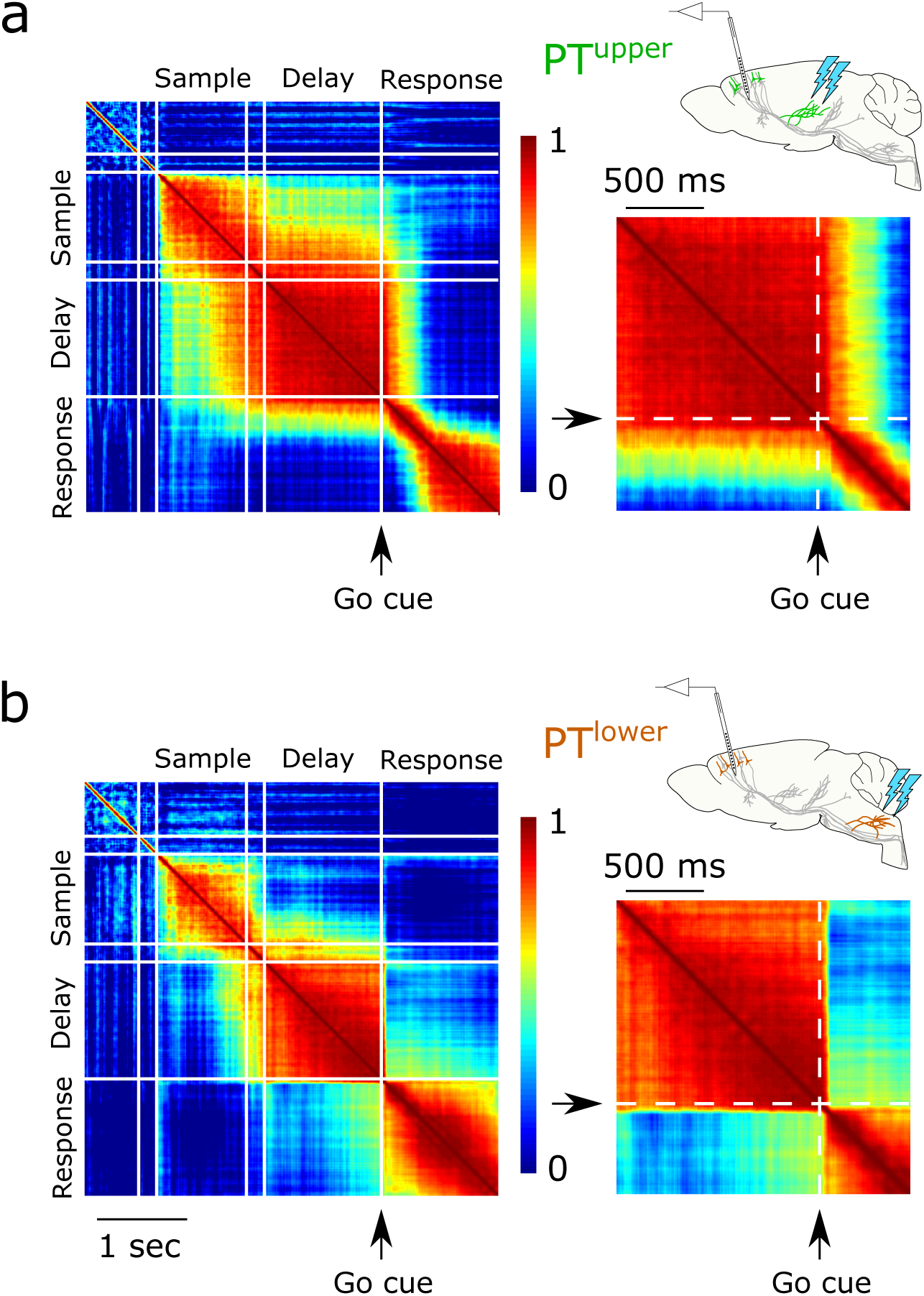
Stability of the coding dimension (CD) across time within a trial. The coding dimension is the dimension that best discriminates trial types in a given time interval. Heat maps represent the correlation of the CD across all pairs of time points. **a.** In PT^upper^ neurons the coding dimension remains similar to CD_early_ for all time points preceding the go cue (including the delay epoch). **b.** In PT^lower^ neurons, the coding dimension in the delay epoch is largely orthogonal to the coding dimension in the sample epoch. Panels (a) and (b) show that the upcoming movement direction is encoded in a persistent manner in the PT^upper^ population, but not the PT^lower^ population. *Right panels:* Expanded view of the change in coding dimension around the time of the go cue. An abrupt change in the coding dimension occurs immediately after the go cue onset in the PT^lower^ population. A change also occurs in the PT^upper^ population, but more slowly (several hundred milliseconds), largely after initiation of movement.

**Extended data figure 11.**
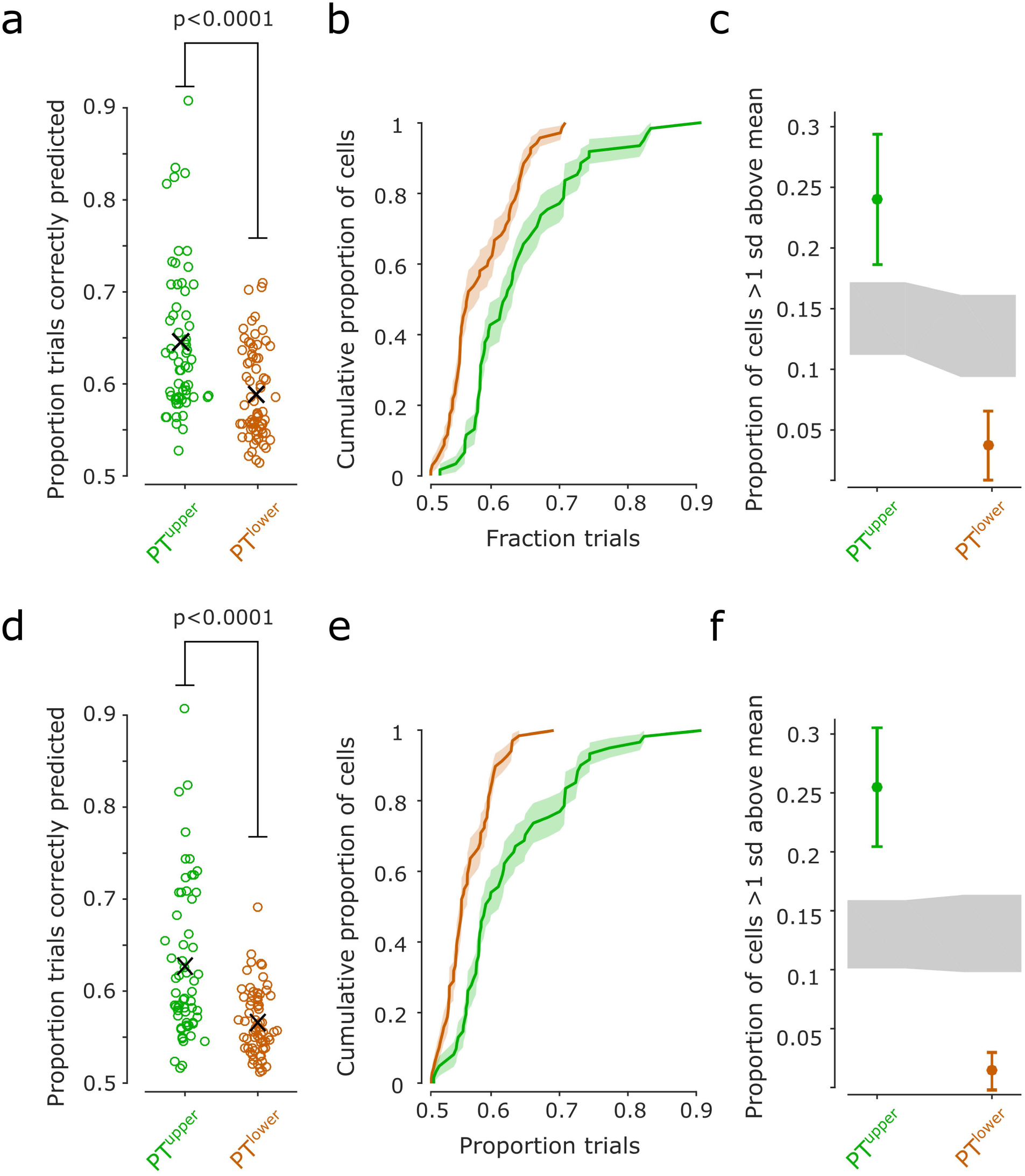
Decoding of trial type in PT neuron types. **a.** Accuracy of trial type classification by single neurons in the 400 ms immediately following stimulus onset. 24.6% (15/61) of PT^upper^ neurons predicted trial type with at least 70% accuracy, whereas only 4.4% (3/69) of PT^lower^ neurons did so. Mean accuracy was also significantly higher in PT^upper^ neurons (PT^upper^: 64.4 ± 1.0%; PT^lower^: 58.9 ± 0.6%, mean ± s.e.; p<0.0001, two-sided Mann-Whitney U-test). **b.** Cumulative distribution function of the data in (a). **c.** Neurons containing the most trial-type information after stimulus onset disproportionately belong to the PT^upper^ class. The 10 most discriminative neurons all belonged to the PT^upper^ population. **d-f.** Same as (a-c) but decoding only based on spike rate increases above baseline. Trial-type selectivity during the sample epoch in PT^lower^ neurons was predominantly characterized by a modest suppression of spiking on one trial type, likely reflecting widespread lateral inhibition. Disregarding spike rate changes below baseline, no PT^lower^ neurons predicted trial type with at least 70% accuracy, while the same 24.6% of PT^upper^ neurons continued to do so and accounted for 20/21 of the most predictive neurons (PT^upper^: 62.7 ± 1.1%; PT^lower^: 56.7 ± 0.4%, mean ± s.e.; p<0.0001, two-sided Mann-Whitney U-test). As soon as the trial type is cued by the stimulus, upcoming movement direction is encoded robustly in the PT^upper^ population and only minimally in PT^lower^ cells.

**Extended data figure 12.**
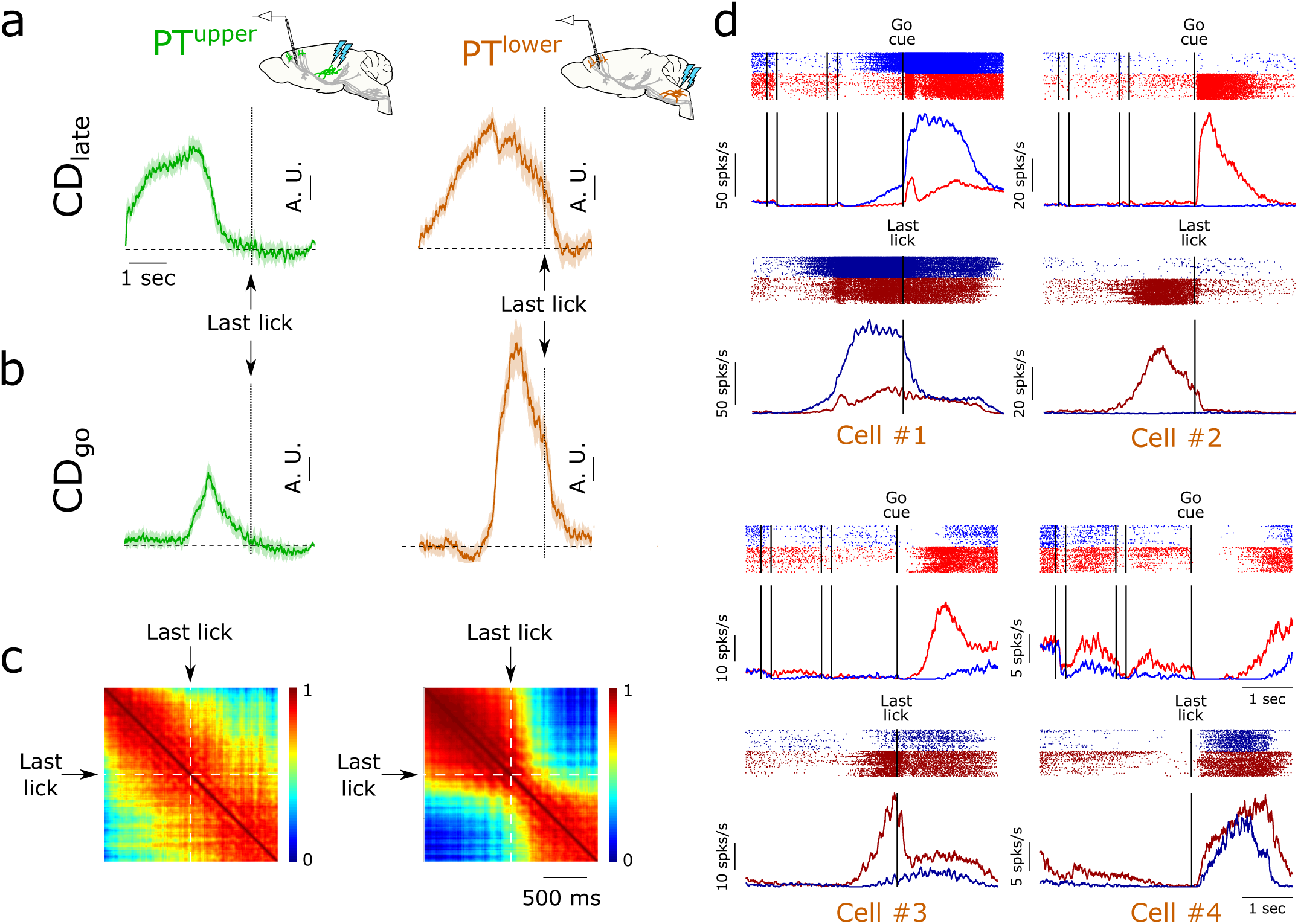
Movement termination signals in PT^lower^ neurons. **a.** Selectivity along CD_late_ (same as **EDFig. 9**) for PT^upper^ (green; *left*) and PT^lower^ neurons (orange; *right)* aligned to the last lick in the response epoch. **b.** Selectivity along CD_go_ (same as **Fig. 6**) aligned to the last lick for each PT type. **c.** Correlation of coding dimension weights at all pairs of time points after the go cue for PT^upper^ neurons (*left*) and PT^lower^ neurons (*right*) using last-lick aligned spike rates. An additional transition in the population dynamics accompanies the termination of movement in PT^lower^ neurons, while there is no correlate of movement termination in PT^upper^ neurons. The change in dynamics at the offset of movement was somewhat less abrupt than at movement onset, likely a result of aligning data to the last lick port contact, which does not precisely mark the cessation of movement. **d.** Spike raster plots (*top*) and trial-averaged activity (*bottom*) for four example PT^lower^ neurons aligned to the go cue (lick right: *blue;* lick left: *red*) and the last lick port contact (lick right: *dark blue;* lick left: *dark red*).

## EXTENDED DATA TABLES

**Extended data table 1.** Digital object identifiers for reconstructed PT neurons.

| Type | Neuron ID | DOI | Complete? |
| --- | --- | --- | --- |
| Thalamus-projecting (PT<sup>upper</sup>) | AA0011 | https://doi.org/10.25378/janelia.5521615 | Y |
| Thalamus-projecting (PT<sup>upper</sup>) | AA0114 | https://doi.org/10.25378/janelia.5526721 | Y |
| Thalamus-projecting (PT<sup>upper</sup>) | AA0115 | https://doi.org/10.25378/janelia.5526724 | Y |
| Thalamus-projecting (PT<sup>upper</sup>) | AA0122 | https://doi.org/10.25378/janelia.5527240 | Y |
| Thalamus-projecting (PT<sup>upper</sup>) | AA0181 | https://doi.org/10.25378/janelia.5527444 | Y |
| Thalamus-projecting (PT<sup>upper</sup>) | AA0182 | https://doi.org/10.25378/janelia.5527447 | Y |
| Thalamus-projecting (PT<sup>upper</sup>) | AA0245 | https://doi.org/10.25378/janelia.5527657 | Y |
| Thalamus-projecting (PT<sup>upper</sup>) | AA0261 | https://doi.org/10.25378/janelia.5527717 | Y |
| Medulla-projecting (PT<sup>lower</sup>) | AA0012 | https://doi.org/10.25378/janelia.5521618 | Y |
| Medulla-projecting (PT<sup>lower</sup>) | AA0133 | https://doi.org/10.25378/janelia.5527273 | Y |
| Medulla-projecting (PT<sup>lower</sup>) | AA0179 | https://doi.org/10.25378/janelia.5527438 | Y |
| Medulla-projecting (PT<sup>lower</sup>) | AA0180 | https://doi.org/10.25378/janelia.5527441 | Y |
| Thalamus-projecting (PT<sup>upper</sup>) | AA0131 | https://doi.org/10.25378/janelia.5527267 | N |
| Thalamus-projecting (PT<sup>upper</sup>) | AA0134 | https://doi.org/10.25378/janelia.5527276 | N |
| Thalamus-projecting (PT<sup>upper</sup>) | AA0169 | https://doi.org/10.25378/janelia.5527408 | N |
| Medulla-projecting (PT<sup>lower</sup>) | AA0132 | https://doi.org/10.25378/janelia.5527270 | N |
| Medulla-projecting (PT<sup>lower</sup>) | AA0135 | https://doi.org/10.25378/janelia.5527279 | N |
| Medulla-projecting (PT<sup>lower</sup>) | AA0250 | https://doi.org/10.25378/janelia.5527678 | N |

**Extended data table 2.** Experiments

| Figure panel(s) | Animals (group #) | Viruses injections (#) | Notes |
| --- | --- | --- | --- |
| Fig. 2 c,d and EDFig. 4 | 1 | 1,2,3,4 | Retrograde nuclear labeling from medulla, thal., SC, pons |
| EDFig. 3 | 2 | 5,6 | Retrograde labeling from medulla and thal. |
| Fig. 3b,c | 3 | 7,8 | Retrograde labeling from medulla and thal. |
| Fig 4-6, EDFig. 3 and 7-12 | 4,5 | 9,10,11 | Cell-type specific electrophysiology |
| Fig. 6 | 6 | N/A | Reaction time measurement |
| Fig. 2a-b, EDFig. 2 | 7 | 12,13 | Axonal reconstruction |
| EDFig. 5,6 | 8 | 14,15 | Retrograde labeling from medulla and thal. For bulk RNA-Seq |

**Extended data table 3.** Animals used in experiments

| Group number | Genotype | Source | Catalog # | Cohort |
| --- | --- | --- | --- | --- |
| 1 | C57BL/6 | Charles River | 027 | 2F |
| 2 | EMX1-IRES-Cre | JAX | 005628 | 3M |
| 3 | EMX1-IRES-Cre | JAX | 005628 | 4M |
| 4 | C57BL/6J | JAX | 000664 | 3M |
| 5 | EMX1-IRES-Cre | JAX | 005628 | 7M, 2F |
| 6 | EMX1-IRES-Cre | JAX | 005628 | 4M |
| 7 | C57BL/6 | Charles River | 027 | 5F |
| 8 | C57BL/6 | Charles River | 027 | 7F |

**Extended data table 4.** Virus injections

| Injection number | Injection name | Virus name | Source | Plasmid Catalog # | Titer (GC/mL) | Coordinate | Volume (nL) |
| --- | --- | --- | --- | --- | --- | --- | --- |
| 1 | Medulla tdT (nuclear) | AAV2-retro-CAG-H2B-tdTomato | JRC |  | 1.3e13 | M/L 1.25 (left), A/P -6.6 (bregma), D/V 4.1 and 4.5 | 40 |
| 2 | Thalamus GFP (nuclear) | AAV2-retro-CAG-H2B-GFP | JRC |  | 1.2e13 | M/L 1.0 (right), A/P -1.5 (bregma), D/V 3.2 | 40 |
| 3 | SC tdT (nuclear) | AAV2-retro-CAG-H2B-tdTomato | JRC |  | 1.3e13 | M/L 1.1 (right), A/P +0.3 (lambda), D/V 2.2 | 40 |
| 4 | Pons FLAG (nuclear) | AAV2-retro-CAG-H2B-mRuby2-smFP FLAG | JRC |  | 6.7e12 | M/L 0.4 (right), A/P +0.1 (lambda), D/V 5.5 | 40 |
| 5 | Medulla tdT | AAV2-retro-CAG-tdTomato | JRC |  | 5.5e12 | M/L 1.25 (left), A/P -6.6 (bregma), D/V 4.1 and 4.5 | 20 (x2) |
| 6 | Thalamus GFP | AAV2-retro-CAG-GFP | JRC | Addgene# 28014 | 1.3e13 | M/L 1.0 (right), A/P -1.5 (bregma), D/V 3.2 | 30 |
| 7 | Medulla FLAG | AAV2-retro-CAG-mRuby2-smFP FLAG | JRC | Addgene# 59760 | 9.5e12 | M/L 1.25 (left), A/P -6.6 (bregma), D/V 4.1 and 4.5 | 20 (x2) |
| 8 | Thalamus FLAG | AAV2-retro-CAG-mRuby2-smFP FLAG | JRC | Addgene# 59760 | 9.5e12 | M/L 1.0 (right), A/P -1.5 (bregma), D/V 3.2 | 40 |
| 9 | Medulla Chr2 | AAV2-retro-Syn-ChR2(H134R)-GFP | JRC |  | 2.0e13 | M/L 1.25 (left), A/P -6.6 (bregma), D/V 4.3 | 200 |
| 10 | Medulla Flex Chr2 | AAV-SL1-Syn-Flex-ChR2-YFP | JRC |  | 1.9e13 | M/L 1.25 (left), A/P -6.6 (bregma), D/V 4.3 | 60 |
| 11 | Thalamus Flex Chr2 | AAV-SL1-Syn-Flex-ChR2-YFP | JRC |  | 1.9e13 | M/L 1.0 (right), A/P -1.7 (bregma), D/V 3.2 | 50 |
| 12 | ALM sparse tdT | AAV2/1-Syn-iCre<br>AAV2/1-CAG flex rev-tdTomato | JRC |  | 2.1e8<br>9.0e12 | M/L 1.2 (left), A/P +2.3 (bregma), D/V 0.4 and 0.9 | 20 (x2) |
| 13 | ALM sparse GFP | AAV2/1-Syn-iCre<br>AAV2/1-CAG flex rev-3xGFP | JRC |  | 2.1e8<br>1.4e13 | M/L 1.2 (left), A/P +2.3 (bregma), D/V 0.4 and 0.9 | 20 (x2) |
| 14 | Medulla tdT | AAV2-retro-CAG-tdTomato | JRC |  | 5.5e12 | M/L 1.25 (left), A/P -6.6 (bregma), D/V 4.1 and 4.5 | 20 (x2) |
| 15 | Thalamus GFP | AAV2-retro-CAG-GFP | JRC | Addgene# 28014 | 1.3e13 | M/L 1.0 (right), A/P -1.5 (bregma), D/V 3.2 | 20 |

